# Experimentally revealed stochastic preferences for multi-component choice options

**DOI:** 10.1101/832881

**Authors:** Alexandre Pastor-Bernier, Konstantin Volkmann, Arkadiusz Stasiak, Fabian Grabenhorst, Wolfram Schultz

## Abstract

Realistic, everyday rewards contain multiple components. An apple has taste and size. However, we choose in single dimensions, simply preferring some apples to others. How can such single-dimensional preference relationships refer to multi-component choice options? Here, we measured how stochastic choices revealed preferences for two-component milkshakes. The preferences were intuitively graphed as indifference curves that represented the orderly integration of the two components as trade-off: parts of one component were given up for obtaining one additional unit of the other component without a change in preference. The well-ordered, non-overlapping curves satisfied leave-one-out tests, followed predictions by machine learning decoders and correlated with single-dimensional Becker-DeGroot-Marschak (BDM) auction-like bids for the two-component rewards. This accuracy suggests a decision process that integrates multiple reward components into single-dimensional estimates in a systematic fashion. In inter-species comparisons, human performance matched that of highly experienced laboratory monkeys, as measured by accuracy of the critical trade-off between bundle components. These data describe the nature of choices of multi-component choice options and attest to the validity of the rigorous economic concepts and their convenient graphic schemes for explaining choices of human and non-human primates. The results encourage formal behavioral and neural investigations of normal, irrational and pathological economic choices.

## Introduction

We like sweet apples. We can state our preference in words, but they may not be accurate because of poor introspection, faulty memory or erroneous report. It would be better to observe our choice. But how do we choose apples? We may prefer a sweeter apple even if it is a bit smaller; hence we trade-in size for sweetness. Our preference does not concern any component alone but their combination. Every reward or economic good has multiple components, attributes or dimensions, and thus constitutes a bundle. The bundle components may be integral parts of a good, like sweetness and size of an apple, or consist of distinct entities, like steak and vegetable of a meal. Each component contributes to the choice. Without considering the multi-component nature of choice options, we would only study exchanges, like choosing between an apple and a pear (not a choice for an apple lover), or between a movie and a meal (not good when hungry). Thus, to understand realistic choice, we should consider the multi-component nature of choice options.

In contrast to the multi-dimensionality of choice options, their subjective value, or utility, varies only along one dimension. Likewise single-dimensional are the preference relationships that are revealed by our choice of bundles. With two options, a rational decision maker prefers either one option, or its alternative, or is indifferent to them (completeness axiom; Von Neumann & Morgenstern, 1944; Mas-Colell, Whinston, & Green, 1995). With repeated stochastic choice, preferences are revealed by choice probability (McFadden & Richter, 1990; McFadden, 2004); the probability of choosing one option over its alternative varies in a graded, scalar manner. Correspondingly, the utility of a choice option can only be higher, lower or equal to that of its alternative. Further, neural signals representing choice options can only vary along a single dimension at any given moment and thus are also scalar; their firing rate either increases, remains unchanged, or decreases (thus constituting a distribution), even when encoding multi-dimensional variables (Pastor-Bernier, Stasiak, & Schultz, 2019). Hence the question: how can single-dimensional preferences, utility and neural signals concern multi-component choice options? Or, put more formally, how can scalar measures reflect vectorial bundles? And can we test the issue empirically, using well worked-out, rigorous theoretical concepts that should reduce possible confounds?

### Concepts and hypotheses

The issue of vectorial-to-scalar transformation of economic goods during economic decisions can be addressed by studying revealed preferences. The key notion posits that preferences cannot be observed directly but are revealed by measurable choice. Revealed preferences contrast with stated preferences; both are unobservable, but only revealed preferences refer to immediate, actual choices. Revealed preferences may be inherent and fixed, sitting there and waiting to be revealed by the choice. This is a basic assumption of formal Revealed Preference Theory (Fisher, 1892; Samuelson, 1937; Samuelson, 1938). However, preferences vary with many factors such as context, frame and number of options, which might indicate that preferences are flexible and constructed on the fly, at the moment of choice. These distinctions are important and a matter of on-going debate (Payne, Bettman, & Schkade, 1999; Simonson, 2008; Dhar & Novemsky, 2008; Kivetz, Netzer & Schrift, 2008; Warren, McGraw & Van Boven, 2011). However, the possibility to empirically infer preferences from observable choice is invaluable for investigating choices of multi-component options. Therefore, we like to restrict the use of the term ‘revealed’ to the inference of preferences from observable choice irrespective of their assumed origin, while nevertheless benefiting from the rigorous concepts and graphics of Revealed Preference Theory.

We used the following notions of Revealed Preference Theory and stochastic choice theories (Luce 1959; McFadden & Richter, 1990; McFadden, 2004; Stott, 2006) for humans (for the design of our parallel study on monkeys, see Method; Pastor-Bernier, Plott, & Schultz, 2017):

1. The option set contains two simultaneously presented bundles (binary choice). Each bundle contains two components (A and B). The options are mutually exclusive (choose one bundle or its alternative but not both) and collectively exhaustive (they contain all available bundles).
2. Each bundle can be plotted at the intersection of an x-coordinate (component A) and an y-coordinate (component B) on a two-dimensional graph (Figure 1A).
3. Stochastic preference is revealed by the probability of choosing one bundle over its alternative, which depends on all components of both bundles.
4. Every bundle has a utility for the decision maker that depends only on the amounts of both of its components. A bundle is chosen with higher probability than any other bundle in the same option set if and only if its utility is higher than in any other bundle in that option set. In other words, the preference relationship between two bundles is monotone if the higher utility of one bundle implies that it is preferred to its alternative (Mas-Colell, Whinston, & Green, 1995).
5. Two bundles are chosen with equal probability if and only if their utility is the same. Equal choice probability of *P* = 0.5 for each bundle reveals equal stochastic preference and is graphically represented by a two-dimensional indifference point (IP). At the IP, some amount of one component is given up in order to gain one unit of the other component without change in preference (marginal rate of substitution, MRS) (Figure 1A). In the apple example, some size is given up for more sweetness (the sweeter apple was smaller). The IP is estimated from an S-shaped psychophysical function fitted to the choice probabilities while varying one component of one bundle and keeping all other components constant (Figure 1C, D). The trade-off at the IP is conceptually important, as it demonstrates same preference despite oppositely varying bundle composition.
6. Multiple IPs align on a single, continuous indifference curve (IC). The ICs are graphically characterized by two parameters: (i) the slope, which reflects the relative utility (currency) of the two bundle components; it could be asymmetric between x-axis and y-axis; (ii) the curvature, which captures any slope change between IC center and IC periphery. The ICs are typically convex (viewed from the origin): larger reward amounts are associated with decreasing value increment (reflecting diminishing marginal utility of concave utility function) and therefore require larger trade-in amounts for getting a smaller amount of the other component.
7. Bundles with larger amounts of one or both components are plotted farther away from the origin and are assumed to be revealed preferred to bundles with smaller amounts (the ‘value function’ for the component is strictly monotonically positive; ‘more is always better’). Hence, any bundle above an IC (farther away from origin) would be revealed preferred to any bundle on that IC, and any bundle on an IC would be revealed preferred to any bundle below that IC (closer to origin).
8. The preference relationship between two bundles may hold even when one component of the preferred bundle has a smaller amount than the alternative bundle (partial physical non-dominance, requiring overcompensation by the other bundle component). This aspect, together with the equal-preference trade-off, crucially reflects the integration of the physical value of both bundle components into single-dimensional preference relationships and utility.

An alternative to integrating multiple bundle components may exist when choices follow the amounts of only one bundle component and partly or fully neglect changes in the other component. With such ‘lexicographic’ preferences, the decision maker would forego potential gains from the neglected component and thus violate the principle of utility maximization. In such cases, ICs would be parallel to one of the axes: any bundle on such a line would be equally preferred, and more preferred bundles would lie on one or more parallel lines farther out on the graph. Such scenarios are not far-fetched, as decision-makers may have different discrimination thresholds for different bundle components, such as seen with reward amounts and probabilities (Tversky 1969; Just-Noticeable-Difference, JND; Rieskamp, Busemeyer, & Mellers, 2006).

**Figure 1.**
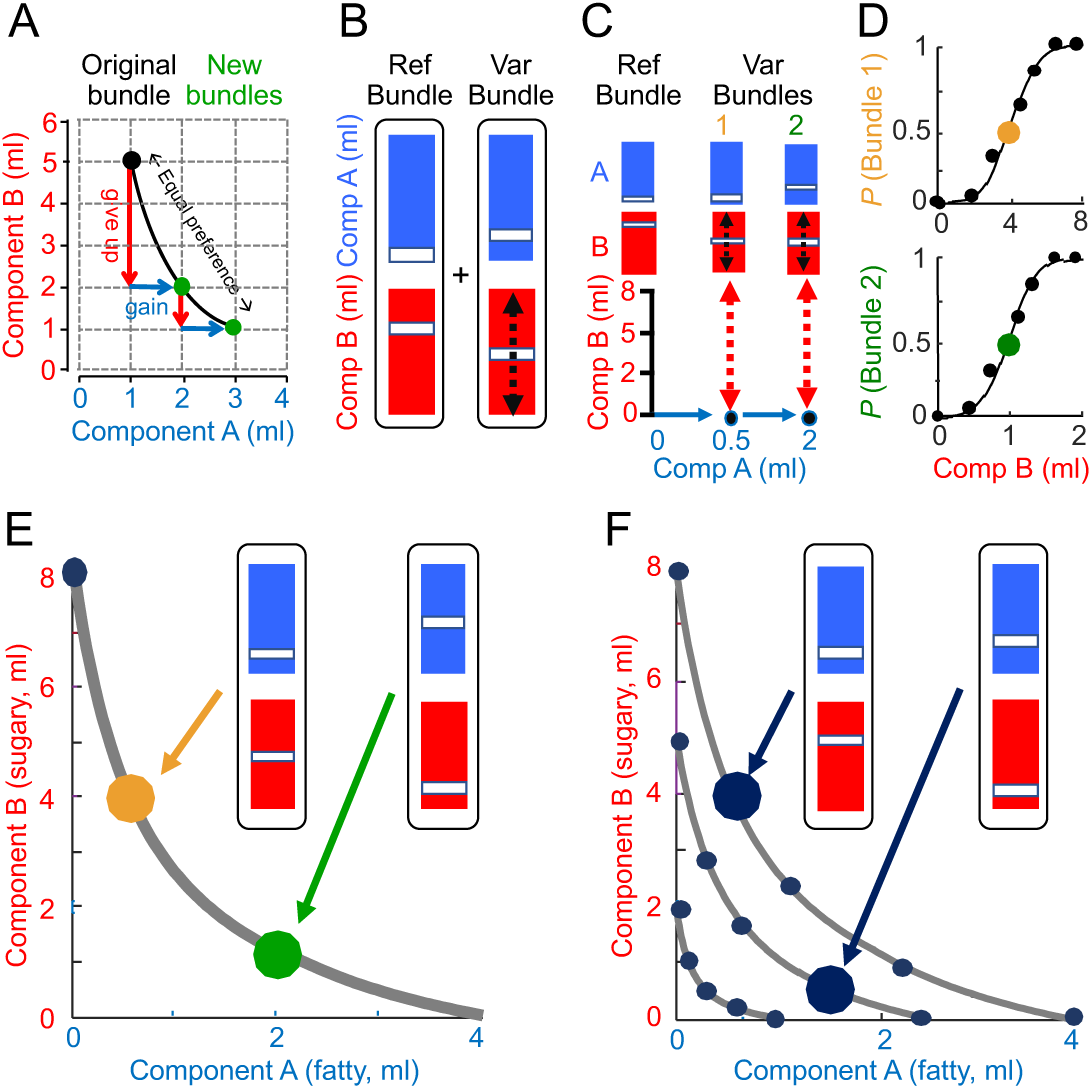
Concepts, binary choice task and experimental design. (A) Scheme of trade-off on indifference curve. Equal revealed preference from varying bundle composition (inspired by a textbook scheme; Perloff, 2009). The original (black) and new bundles (green) are equally revealed preferred (black curve) when a quantity of component B is traded in (red; Marginal Rate of Substitution: initially 3 ml, then 1 ml) for one unit of component A (green; 1 ml). (B) Bundle stimuli for binary choice. Each bundle contained two components with amounts between 0.0 and 8.0 ml. In this example, the Reference Bundle contained a low amount of Component A (Comp A: low-sugar high-fat milkshake) and a high amount of Component B (Comp B: high-sugar low-fat milkshake). Participants chose between the Reference Bundle and the Variable Bundle, whose locations on a computer monitor alternated pseudorandomly between fixed left and right positions. (C) Psychophysical test design. In the Reference Bundle, components A and B were set to 0.0 ml and 8.0 ml, respectively; in the Variable bundle, component A was set to a participant-specific test amount (Table S1A-C) while psychophysically varying the amount of component B (dashed arrows). (D) Psychophysical assessment of two example choice indifference points (IP; choice probability P = 0.5 each option; yellow and green) in a typical participant. During repeated trials, the participant chose between the preset Reference Bundle and the Variable Bundle. Component A of the Variable Bundle was set to pre-determined participant-specific test amounts (here 0.5 ml and 2.0 ml). Component B of the Variable Bundle varied between seven randomly selected amounts of component B. IPs were estimated from choice probabilities, p (Bundle 1) and p (Bundle 2), respectively, using the probit model (Eqs. 1, 2). (E) Schematic indifference curve (IC), fitted by a hyperbola (Eqs. 3, 3a) to all equally revealed preferred but differently composed bundles (‘indifference points’, IPs), as tested in binary choice between an anchor bundle on the y-axis (blue dot, top left) and psychophysically varied test bundles (yellow and green dots), as estimated in D. (F) Schematic map of three hyperbolically fitted ICs. Increasing distance from origin reflects larger milkshake amounts and represents higher utility; all bundles on higher ICs were revealed preferred to all bundles on lower ICs. Arrows denote a preference relationship between two bundles with oppositely varying physical amounts of component A (top-IC bundle is revealed preferred to mid-IC bundle despite lower physical amount of component A; partial physical non-dominance).

### Previous studies

Without referring to the IC graphism, studies tested quality and price of television sets (Simonson, 1989), comfort and fuel consumption of cars (Simonson 1989), payoff amount and probability (Tversky, 1969; Soltani, De Martino, & Camerer, 2012; Levy, Belmaker, Manson, Tymula, & Glimcher, 2012; Chau, Kolling, Hunt, Walton, & Rushworth 2014), shape and color of cards (Fellows 2016), artificial objects and appendages (Pelletier & Fellows, 2019), various food components (Suzuki, Cross, & O’Doherty, 2017; DiFeliceantonio et al., 2018; Harris, Clithero, & Hutcherson, 2018), hats and shoes (Thurstone, 1931), and pastries and payoffs (MacCrimmon & Toda, 1969). Studies using ICs were restricted to hypothetical outcomes, such as hats and shoes (Thurstone, 1931), pencils and payoff (MacCrimmon & Toda, 1969), and monetary token (Choi, Fisman, Gale, & Kariv, 2007; Kurtz-David, Persitz, Webb, & Levy, 2019). Without experimentally establishing ICs, studies used the IC scheme for conveniently explaining choice inconsistencies when an added non-preferred option to the choice set affects the preferences of the original option and thus violates the Independence of Irrelevant Alternatives (IIA) (like adding a reasonably priced decent camera alters the choice between an expensive but fantastic camera and a cheap but bad camera; ‘compromise effect’). Such inconsistencies might be particularly frequent with multi-component bundles that require attention to and integration of the components in addition to the already complex economic decision process itself; difficulties are seen in normal humans (Tversky 1969; Rieskamp, Busemeyer, & Mellers, 2006; Chung et al., 2017; Gluth, Hotaling, & Rieskamp, 2017; Li, Michael, Balaguer, Castanon, & Summerfield, 2018), patients with frontal lobe lesions (Fellows, 2006; Fellows, 2016; Pelletier & Fellows, 2019) and animals (Silberberg, Widholm, Bresler, Fujita, & Anderson 1998; Beran, Ratliff, & Evans, 2009; Pattison & Zentall, 2014; Zentall, 2019). Despite the importance of these economic concepts, what is lacking are more empirical tests of the IC formalisms in controlled laboratory situations using actual, tangible outcomes. Such experiments should help to describe choices involving multi-component options, empirically scrutinize the theoretical concepts and encourage future behavioral and neuroeconomic studies on multi-component choices and their fine-grained ‘irrational’ IIA anomalies in normal and brain-damaged humans who might fail to properly consider all components of choice options.

### The current study

Our experiment revealed stochastic preferences in humans and compared human performance with that of monkeys. Monkeys, as our evolutionary close neighbors, have superb and understandable cognitive and behavioral abilities that allow generalization across primate species and high-resolution neuronal investigations. Inter-species comparisons are also methodologically useful, as monkeys can participate longer in experiments and provide much larger numbers of trials for more sophisticated tests, including preference between bundles not used for IC fitting, transitivity across partially physically non-dominating bundles on different ICs, and change of option set size, all in the same individual.

The perspective of serving as template for future human and animal studies determined the experimental design and imposed constraints concerning event timing, trial repetition and immediately consumable payouts comparable to tangible liquid and food rewards for animals (Kagel et al., 1975). We followed the notions above and used the same design and data analysis as in rhesus monkeys (Pastor-Bernier, Plott, & Schultz, 2017). Participants repeatedly chose between two composite visual stimuli presented on a computer monitor. Each stimulus predicted a bundle of the same two milkshakes that varied in amount of fat and sugar concentration and were delivered directly to the participant’s mouth (component A, component B; Figure 1B). The milkshakes were fully known to the participants and constituted rewards, as evidenced by the participants’ voluntary consumption. We kept effort cost constant and equal for both choice options, but we did not test budget constraints to stay compatible with neurophysiological primate studies in which budgets would add confounds for interpreting neuronal responses and thus increase experimental complexity. We estimated ICs from psychophysically assessed IPs and confirmed the resulting two-dimensional preference map with mechanism-independent Becker-DeGroot-Marschak (BDM) auction-like bidding that truthfully reveals the bidder’s value of desired objects in humans and monkeys (Becker, DeGroot, & Marschak, 1964; Plassmann, O’Doherty, & Rangel, 2007; Al-Mohammad & Schultz, 2019). The choices conformed with the ICs of economic theory, satisfied leave-out statistics, corresponded to decoder predictions, and correlated with BDM bids. The trade-off accuracy of humans compared well with that of laboratory monkeys.

## Method

### Human participants

A total of 24 human participants (11 males, 13 females; mean age 25.4 years, range 19-36 years) completed a binary choice task for measuring revealed preferences and performed a Becker-DeGroot-Marschak (BDM) control task. The sample size of 24 participants was chosen with future neuroimaging tests on these participants in mind; an earlier study in our laboratory suggested that this sample size was adequate for avoiding false negatives (Zangemeister, Grabenhorst, & Schultz, 2016). The participants were fully informed about the upcoming consumption of milkshakes, the general nature of the binary choice task, the requirement to perform this task and a BDM bidding task in a separate session in the human imaging scanner (whose neural results will be published elsewhere), the remuneration (20 UK £, paid into their bank account a few days after the second session), and the duration of each of the two experimental sessions. Due to these informations, the participants were able to construct the ‘meal’ of milkshakes in advance. None of the participants had diabetes or lactose intolerance, nor did they require specific diets, to avoid medical and cultural interference. All participants had a known appetite for milkshakes and provided written informed consent based on a detailed information sheet. We used similar design, data analysis and validation procedures as in our parallel study on rhesus monkeys, where they are described and discussed in more detail (Pastor-Bernier, Plott, & Schultz, 2017). The Local Research Ethics Committee of the Cambridgeshire Health Authority approved the study.

### Stimuli, rewards and delivery apparatus

The human participants viewed quantitative, colored visual stimuli on a computer monitor, which represented the two milkshakes and their amounts in each of the two bundles (Figure 1B, C). Each bundle stimulus consisted of two vertically aligned rectangles. The color of each rectangle indicated the bundle component. The vertical position of a bar in each rectangle indicated the physical amount of each component (higher was more).

After extensive piloting with various liquids and liquidized foods, we found milkshakes with a controlled mix of sugar and fat to give the most reliable behavioral performance. As the milkshakes were delivered individually (see below), drinks containing either only sugar or only fat were deemed to be too unnatural. Thus, in our bundles, component A (top, blue) consisted of a low-sugar high-fat milkshake (25% whole milk, 75% double cream, no sugar), and component B (bottom, red) consisted of a high-sugar low-fat milkshake (10% sugar in skimmed milk). The Psychtoolbox in Matlab (Version R2015b) running on a Dell Windows computer served for stimulus display and recording of behavioral choices.

Participants were tested one at a time and in the absence of other persons in the room; they were seated on a standard, padded office chair at a standard-height desk in a closed, small, window-less experimental room (ca. 3 x 4 m) with artificial light; the room was located in a quiet laboratory area, without specific ambient noise. Besides the desk and chair, the room was empty except for one standard-height table with a large storage carton box underneath that was unrelated to the experiment. The two milkshakes were delivered directly into the participant’s mouth via a custom-made mouthpiece with two single-use pipette tips onto which the participant bit; the mouthpiece was connected to two silicone tubes approved for delivery of food stuffs; the two tubes were respectively attached to two 50-ml syringes driven by two piston pumps (NE-500, New Era Pump Systems Inc; www.syringepump.com). As used before (Zangemeister, Grabenhorst, & Schultz, 2016), each pump delivered a programmable quantity of one milkshake with milliliter precision (VWR International Ltd) and was controlled by the computer using a National Instruments card (NI-USB-6009) via the Matlab Data Acquisition Toolbox.

### Binary choice task

Each trial started with an initial fixation cross in the center of a computer monitor in front of the participant. After a pseudorandomly varying interval (mean 0.5 s, flat hazard rate from Poisson distribution), the two bundle stimuli appeared simultaneously at pseudorandomly alternating fixed left and right positions on the monitor; each bundle stimulus indicated the same two milkshakes with independently set amounts (Figure 1B). The participant chose one of the two bundle stimuli by pressing a single button once (left or right computer keyboard arrow for corresponding left or right bundle choice), upon which a green rectangle appeared for 200 ms around the chosen bundle to confirm the choice. There was no time limit on the button press, and most reaction times (from appearance of bundle stimuli to button press) were between 0.5 s and 4.0 s (medians of 1.98 s to 2.58 s depending on distance to choice indifference). At 4.0 s after trial start or at 0.5 s after button press, whatever occurred later, all stimuli extinguished, and the participant received either no payout (80% of trials), or a payout (20% of trials); thus, every fifth chosen bundle was paid out on average, using a Poisson distribution. No-payout trials ended here, and a new trial started after an inter-trial interval (ITI) of 0.5 s. Thus, given that button press was allowed to occur later than 4.0 s after trial start, the median duration of unrewarded trials was 5.0 s, and total cycle time (trial + ITI) was 5.5 s.

By contrast, in payout trials, both milkshakes of the chosen bundle were delivered; component A immediately after the choice, and component B at a constant interval of 0.5 s following onset of delivery of component A. The constant delay between the two components, rather than simultaneous delivery or pseudorandomly alternating sequential liquid delivery, served to clearly demarcate the two distinct components and maintain their discriminability. Thus, the utility for component B reflected the subjective value of the milkshake itself and a temporal discount due to longer delay. Delivery of each milkshake by the syringe pump system took between 0.5 s and 5.0 s depending on amount (up to 8.0 ml); thus, the participant could swallow milkshake A while waiting for milkshake B or mix the milkshakes in the mouth before full swallowing. The trial ended 5.0 s following component B delivery onset, which started 0.5 s following component A delivery onset. Adding these 5.5 s for milkshake delivery to the median trial duration of 5.0 s before payout, payout trials lasted about 10.5 s, and total cycle time (trial + ITI of 0.5 s) was 11.0 s.

The full assessment of 3 ICs with 4 IPs required a total of 504 trails (see below for details). With 20% rewarded trials lasting 11.0 s and 80% unrewarded trials lasting 5.5 s, the total duration of an experiment for each participant was 55 min (0.2 x 504 x 11 s + 0.8 x 504 x 5.5 s = 1,109 s + 2,218 s = 3,327 s = 55 min).

Although participants were instructed to not eat or drink up to four hours prior to testing, satiety was a concern due to the high sugar and fat content of the milkshakes. We addressed the issue by the 20% payout schedule, by limiting each payout to maximally 10.0 ml, and by delivering not more than 200 ml of total liquid to each participant on each session. We did not find evidence for satiety with a specific data analysis (Figure 3).

**Figure 2.**
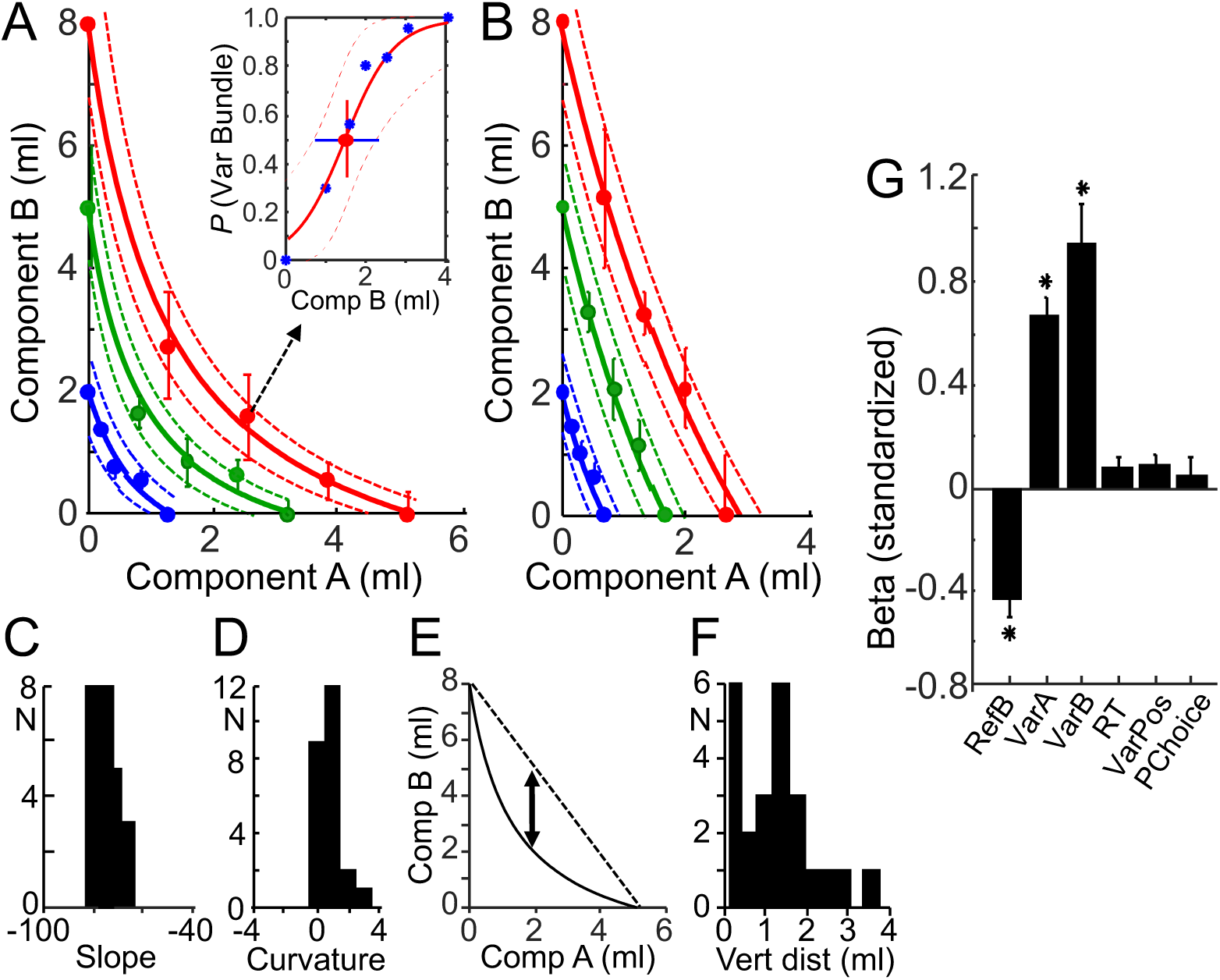
Empirical indifference curves (IC) representing revealed preferences. (A) Typical convex ICs from an example participant, as seen in 18 of the 24 participants. Component A was a low-sugar high-fat milkshake; component B was a high-sugar low-fat milkshake. Solid lines show hyperbolically fitted ICs, dotted lines show 95% confidence intervals of fits. Dots show bundles that are equally preferred on the same IC (IPs). Inset: psychophysical assessment of indifference point (IP) marked on highest IC (test points in blue, IP estimated by probit regression in red). (B) Typical linear ICs from another example participant, as seen in 6 of the 24 participants. (C, D) Distributions of slope and curvature, respectively, of hyperbolically fitted ICs from all 24 participants (coefficients β_2_ / β_1_ and β_3_ in Eq. 3, respectively). N = number of participants. (E) Scheme of intuitive numeric assessment of IC curvature: maximal vertical distance (ml of component B on y-axis) between fitted IC (curve) and a straight line connecting the x-axis and y-axis intercepts. A distance of > 0.0 ml indicates convexity, whereas a 0.0 ml distance indicates perfect linearity. (F) Distribution of convex curvature, as measured using the scheme shown in E. The two peaks indicate six participants each with similarly near-linear ICs and similarly convex ICs, respectively. (G) Specificity of bundle choice, as opposed to unrelated parameters. Bar graph shows standardised beta (β) regression coefficients for choice of Variable Bundle over Reference Bundle (Eq. 5), as assessed for each individual participant and then averaged across all 24 participants. RefB, component B in Reference Bundle; VarA and VarB, components A and B in Variable Bundle; RT, reaction time; VarPos, left-right position of Variable Bundle stimulus; PChoice, choice in previous trial. Error bars show standard error of the mean (SEM). \**P* ≤ 0.02.

**Figure 3.**
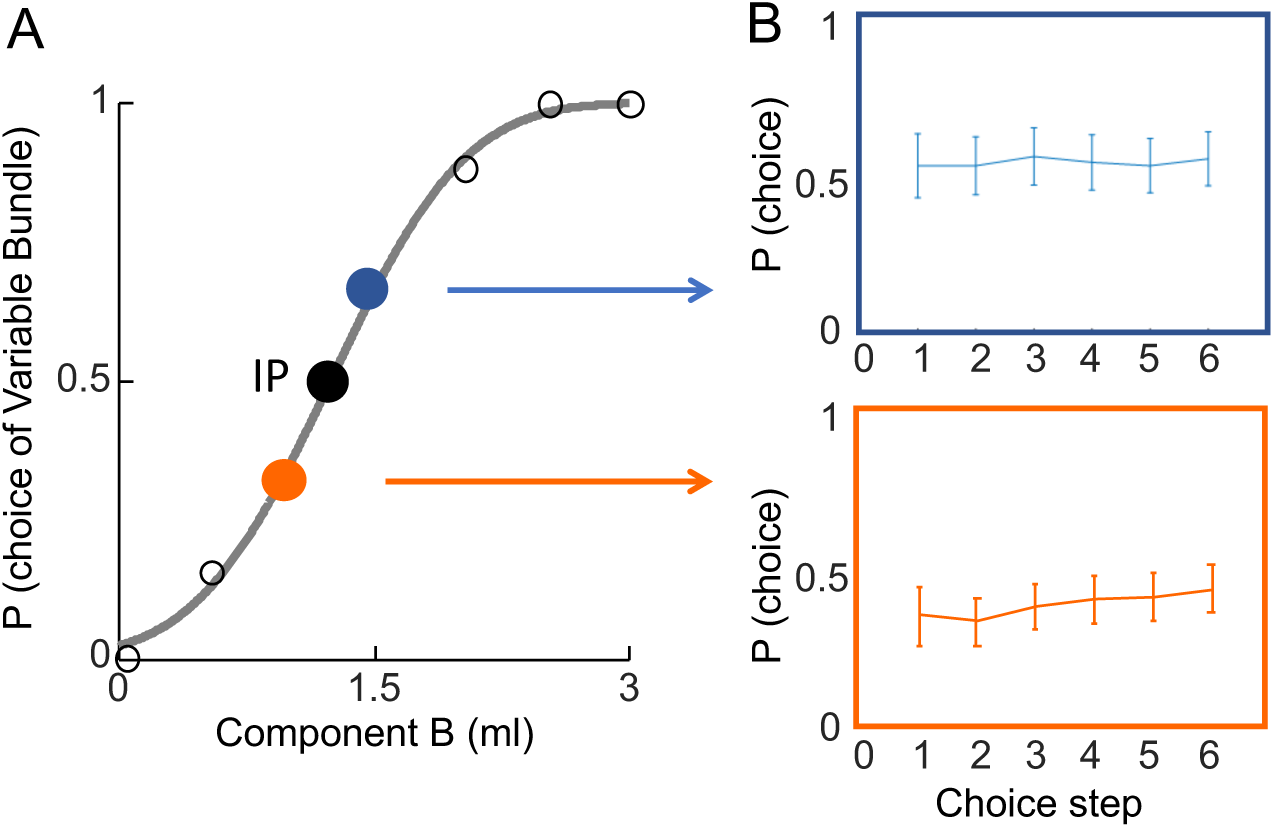
Satiety control. (A) Choice function obtained by sigmoid fit with the probit model. From seven randomly selected fixed amounts of component B in the Variable Bundle, the two colored dots show two amounts associated with choice probabilities closest to the indifference point (IP; p = 0.5 each option). (B) Lack of satiety across typical test duration. Choice probabilities varied insignificantly above (blue) and below (orange) 4 IPs on each of 3 ICs (total of 12 choices tested above and 12 choices tested below IP at each of 6 steps, amounting to 144 choices / participant). Data are averaged from all 24 participants. Total duration of the 6 steps was 20 ± 1.38 min (mean ± SEM).

### Psychophysical assessment of binary choice

We used the same standard psychophysical staircase procedure (Green & Swets, 1966) as in our parallel study on rhesus monkeys (Pastor-Bernier, Plott, & Schultz, 2017) to estimate IPs at which each of two bundles were chosen with equal probability (*P =* 0.5 each option), revealing equal preference for each option. We tested seven bundles for obtaining each IP and, to avoid hysteresis, alternated their sequence randomly. We obtained five IPs for estimating each of three ICs. The procedure required repeated testing, which was also necessary for the subsequent neuroimaging experiment with the same participants (to be reported elsewhere).

We advanced in several steps. First, we realized that participants had different sensitivity ranges for milkshake amounts. To assess the appropriate ranges of good discrimination of both milkshake components, we had each participant chose between single-component bundles containing only milkshake B with the amount of 4 ml (the starting amount for the intermediate IC to be estimated in the full procedure) and amounts varying between 0 and 8 ml of only milkshake A. Once these individual ranges had been determined, the Matlab function *interp1q* was used to set test amounts at five even incremental steps for component A (x-axis) within the specific sensitive range of each participant. The used test amounts for both component for each of the three IC levels are listed in Tables S1A-C and S2. This initial test was also used to discard data from participants with differences in preferences between the two milkshakes larger than 1:3, which would have resulted in technical difficulties for milkshake delivery and control of satiety.

Once the participant-specific test amounts for component A had been determined, we performed the full psychophysical procedure. We set, in the Reference Bundle, component A to 0.0 ml and component B randomly to either 2.0 ml, 5.0 ml or 8.0 ml as starting points for the future three ICs (thus estimating the 3 ICs in random order). Then we set the alternative Variable Bundle as follows: we set the participant-specific test amount of component A one step higher (Step 2 in Table S1A-C), and we randomly selected (without replacement) one of the seven amounts of component B (Table S2); we repeated the selection until all seven amounts had been tested once (Figure 1D). The seven amounts were even divisions of the vertical space (y-axis) between 0 and 2 ml (lowest IC), 0 and 5 ml (intermediate IC), and 0 and 8 ml (highest IC). We performed this procedure six times to estimate each IP using sigmoid fitting (see Eqs. 1, 2 below), thus requiring 42 choices per IP (Figure 1D). At each IP, the amount of component B was usually lower in the Variable Bundle compared to the Reference Bundle. In this way, we implemented the marginal rate of substitution (MRS) at each IP that indicated how much of component B the participant was willing to give up in order to gain one unit of component A, always relative to the constant Reference Bundle.

We obtained three further IPs in choices between the constant Reference Bundle and the Variable Bundle whose amounts of component A increased step-wise (Steps 3, 4 and 5 in Tables S1A-C), thus advancing from top left to bottom right on a two-dimensional x-y indifference map. As the distance in physical space of component B between the Reference Bundle and the Variable Bundle increased, the confidence intervals (CI) increased also rightward for IPs. This effect was consistent across all participants and ICs (however, differences in slope and curvature between participants reflected genuine properties of IC and did not result from the method). We are aware that the unidirectional progression of testing may lead to somewhat different IP estimates than testing in the opposite direction or in random sequence (Knetsch, 1989). However, in this initial study, we were primarily interested in the systematic assessment of consistent IPs rather than exploring potential pitfalls.

We obtained three ICs with the three starting amounts of component B in the Reference Bundle (2.0 ml, 5.0 ml, or 8.0 ml). Each IC required 4 psychophysically estimated IPs (from 5 test bundles); thus the three ICs required a total of 12 IP estimations. Each of the 12 IP estimations involved 7 test amounts and was done 6 times, thus requiring a total of 12 x 7 x 6 = 504 choices for each participant.

### Becker-DeGroot-Marschak (BDM) bidding task

The preferences represented by the ICs should reflect the utility of each IP bundle on a given IC (see notions 4 and 5 above). To confirm the correspondence between revealed preference and inferred utility with an independent estimation method, we assessed the utility of each IP bundle with a BDM mechanism that is akin to a second price auction and estimates the true subjective value of the participant for the bundle (Becker, DeGroot, & Marschak, 1964). The BDM estimates for bundles on IPs should increase with higher ICs but vary only insignificantly along the same IC, thus following the two-dimensional pattern of preferences represented by the ICs.

We ran the BDM task in the same 24 participants as a separate task a few days after the binary choice task during neuroimaging in pseudorandomly selected 50% of trials immediately after a binary choice trial. During neuroimaging, the participant was lying in an fMRI scanner in the supine position (instead of sitting on a chair at a desk) and viewed the stimuli on a computer monitor above the head (instead of the monitor in front). The BDM value bids were then compared with the three levels of revealed preference defined by the IPs estimated in the binary choice task (we compared with revealed preference levels rather than fitted ICs to avoid potential inaccuracies from curve fitting). The data obtained from the binary choice task in the scanner confirmed the currently reported data but will not be further used in the current study; the neuroimaging results will be reported separately.

In the BDM, the participant received a fresh monetary endowment (20 UK pence) on each trial. The participant bid for a bundle against a pseudorandomly set computer bid that was retrieved from a normal distribution with replacement; a normal compared to a uniform distribution slightly focusses behavior into the bidding range of our untrained participants without unduly affecting the cost of misbehavior (Lusk, Alexander, & Rousu, 2007), in analogy to the beta distribution used on monkeys (Al-Mohammad & Schultz, 2019). If the participant’s bid was higher than or equal to the computer bid then the participant received both component rewards of the bundle and paid an amount equal to the computer bid. If the participant’s bid was lower than the computer bid, the participant lost the auction, paid nothing and did not receive any bundle reward. The participant was informed about a win or a loss immediately after placing the bid; when winning the bid, the participant received the bundle rewards in the same sequence and frequency (every fifth trial on average) as in the choice task assessing revealed preferences. Although we did not assess in an objective manner whether the participants understood the BDM, the similarity in payout schedule with the binary choice task and the correlation in performance between the two tasks (see Figure 5) suggested a sufficient amount of comprehension for making BDM a valid mechanism.

**Figure 4.**
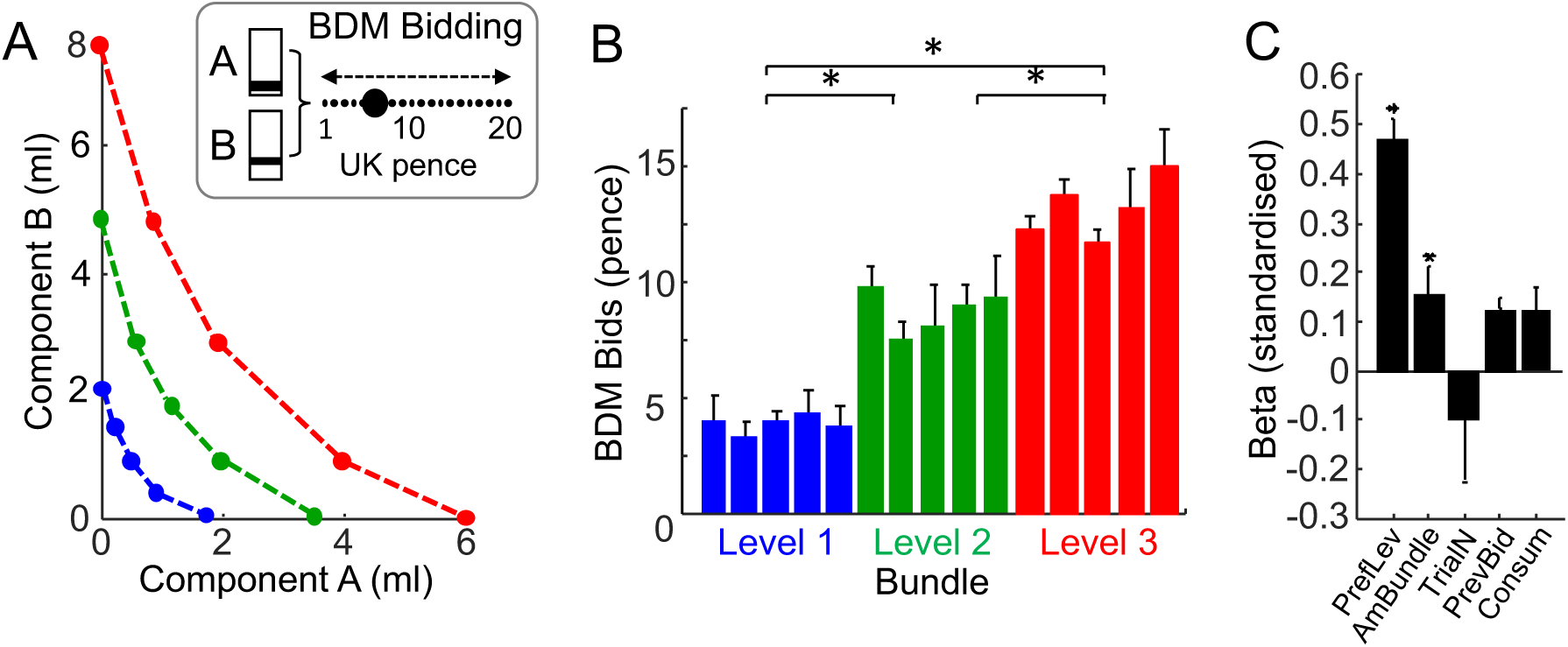
Characteristics of BDM bids for bundles at different revealed preference levels. (A) Schematics of positions of bundles used for eliciting BDM bids at psychophysically estimated points of equal revealed preference (indifference points, IPs, connected by dotted lines). Following the schemes of trade-off and revealed preference, BDM bids should be similar for equally valued bundles (along the dotted lines) but higher for bundles farther away from origin) We tested 5 bundles per level, 3 levels, 12 repetitions, total of 180 bids. Inset: BDM task. Each participant bid for the visually presented two-component (A, B) bundle by moving the black dot cursor using the leftward and rightward horizontal arrows on a computer keyboard. Numbers indicate example bids (in UK pence). (B) Mean BDM bids from a typical participant. The bids were rank-ordered between increasing revealed preference levels (blue, green, red; Spearman Rho = 0.83, P < 0.001) and differed significantly between levels but not within levels (Table S5). Data are shown as mean ± SEM (standard error of the mean), N = 12 bids per bar. (C) Specificity of monetary BDM bids, as opposed to unrelated parameters. Bar graph showing the standardised beta (β) regression coefficients for BDM bids (Eq. 9), as assessed for each individual participant and then averaged across all 24 participants. Abbreviations: PrevLev, revealed preference level (low, medium, high); AmBundle, summed currency-adjusted amount of both bundle components; TrialN, trial number; PrevBid, BDM bid in previous trial; Consum, accumulated drinks consumption. Error bars show SEMs. * P ≤ 0.020.

**Figure 5.**
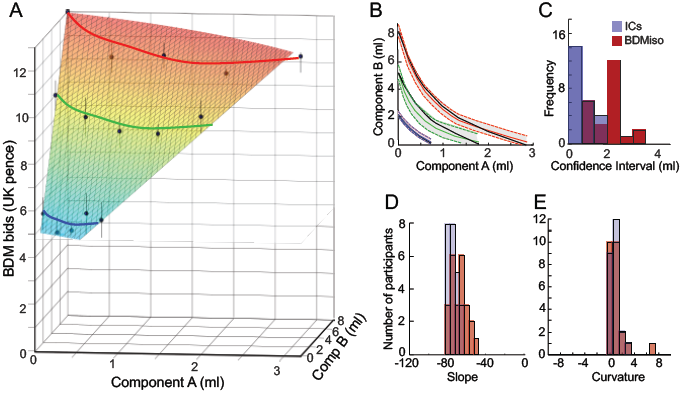
Comparison between BDM bids and indifference curves (IC). (A) Three-dimensional representation of hyperbolically fitted BDM isolines from a typical participant (mean ± SEM). The BDM bids were made for bundles that had been placed on psychophysically estimated revealed preference indifference points (black dots). The BDM isolines were similar along same levels and increased between revealed preference levels. (B) Match between hyperbolically fitted isolines of mean BDM bids (black; Eqs. 9, 9a) and hyperbolically fitted revealed preference ICs (blue, green, red; ± 95% confidence intervals, CI, shaded) from a typical participant. Thus, the BDM isolines fell within the respective CIs of the revealed preference ICs. (C) CIs of fitted BDM isolines (red) and revealed preference ICs (blue) from all 24 participants (averaged along each isoline/IC). The larger 95%CIs of BDM isolines (BDMiso) suggest more variability compared to revealed preference ICs. (D, E) Comparison of slope and curvature coefficient estimates, respectively, between BDM isolines (red) and revealed preference ICs (blue; same data as shown in Figure 2C, D) from all 24 participants.

We showed each participant single bundles that were randomly selected (without replacement) from the set of 15 IP bundles on the 3 revealed preference levels (the same 15 IPs as had been used to fit the 3 ICs shown in Figures 2A, B and S1). A given bundle was set to the participant’s psychophysically estimated IP (Figure 4A). We presented each of the 15 bundles 12 times, resulting in 180 trials in total, and considered the mean of these bids as the BDM-estimated utility. The participant indicated the bid by moving a cursor horizontally on the computer monitor with left and right keyboard arrows (Figure 4A inset). The BDM bid was registered from the cursor position at 5.0 s after onset of presentation of the horizontal bidding scale.

### Statistical analysis

This analysis allowed us to construct the graphic ICs that represent revealed preferences for two-component choice options. Specifically, we used a probit function to estimate each IP bundle from the psychophysical procedure described above, and we employed a hyperbolic regression to obtain each of the three ICs from the respective sets of one start bundle (on the y-axis) and four IP estimations. A separate random effects analysis confirmed that the measured choices were explained by bundle components rather than other regressors.

In order to estimate the IPs numerically, we obtained a sigmoid fit to the empirically assessed choice frequencies via a general linear regression. To do so, we used the Matlab function *glmfit* (Matlab R2015b; Mathworks) on a binomial distribution with a probit link function, which is the inverse of the normal cumulative distribution function (G). Specifically, the generalised linear regression y = β_0_ + β_1_B_var_ + ε can be rewritten after applying the link function as:

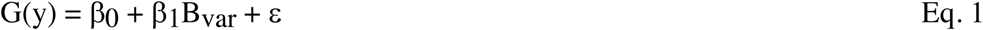

with y as number of times the subject chose the Variable Bundle in the current block from a series of six repetitions, β_0_ as offset constant, β_1_ as regression slope coefficient, B_var_ as reward amount (ml) of component B in the Variable Bundle, and ε as residual error. We chose the probit model because it assumes that random errors have a multivariate normal distribution, which makes it attractive as the normal distribution provides a good approximation to many other distributions. The model does not rely on the assumption of error independence and is used frequently by econometricians (Razzaghi, 2013). Further, preliminary data analysis had revealed a slightly better fit for the probit model compared to the logit model (deviance of 0.4623, as twice log-likelihood difference between probit model and maximum-parameter model, compared to deviance for logit model of 0.5907; 3,200 trials, 5 participants). Therefore we estimated the IPs from the sigmoid fit provided by the probit model, using the following equation:

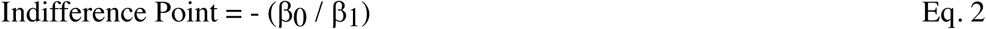

with β_0_ and β_1_ as coefficients of the general linear regression (Eq. 1).

We obtained single ICs, separately for each individual participant, from a set of individual IPs by weighted least-mean-square, non-linear regression (as opposed to the probit regression for estimating each IP). We applied a weight in order to account for within-participant choice variability; the weight was the inverse of the standard deviation of the titrated amount of the B-component at the corresponding IP (the IP having been estimated by the probit regression). We estimated the best-fitting β coefficients from least-mean-square fitting to obtain the equal-preference IC (which is a utility level) and wrote the basic hyperbolic equation in our notation as:

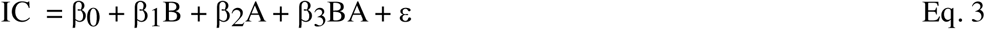

with A and B as amounts of components A and B (ml) referring to the x-axis and y-axis, respectively, β_1_ and β_2_ as slope coefficients of the regressors B and A, and β_3_ as curvature coefficient. The overall slope of the IC itself (global MRS) is calculated as y/x; as components B and A extend on the y-axis and x-axis, respectively, and as β is inversely related to the impact of physical amount of the respective component, the IC slope is (1/β_1_)/(1/β_2_) or β_2_/β_1_.

As IC is a constant, we merged the other constants offset (β_0_) and error (ε) into a common final constant k. To draw the ICs, we computed the amount of component B as a function of component A from the derived equation:

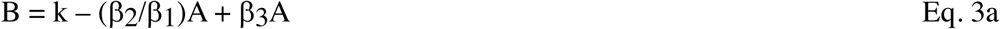

To graphically display a fitted IC (Figure 2A, B), we plotted the preset amount of component A on the x-axis, and the computed fitted amount of component B (Eq. 3a) on the y-axis. The error on the hyperbolic curve was measured as 95% CI. The higher the error around an IP the less weight was given to this point when the IC was calculated. This model resulted in good fits in earlier work (Pastor-Bernier, Plott, & Schultz, 2017). In this way, the IPs of 5 equally revealed preferred but differently composed bundles aligned as a single fitted IC. The three ICs representing increasing revealed preference levels (low, medium, high) were located increasingly farther away from the origin (Figures 1F; 2A, B). The indifference map of 3 × 5 IPs was unique for each participant (Figure S1).

The IC shape was derived from hyperbolic fit and was quantified by two coefficients: slope and curvature. The IC slope coefficient, derived from the ratio of regression slope coefficients (β_2_ / β_1_), reflected the currency relationship between the components and described the participant’s preference for component A relative to component B. For example, an IC slope of - 60°indicated that component A was valued twice as much as the same ml amount of component B. The curvature coefficient (β_3_) quantified the constancy in the trade-off between bundle components. A linear curve (curvature coefficient = 0) indicated a constant rate of exchange for the bundle components, suggesting that the components were perfect substitutes. A more convex IC (curvature coefficient > 0) indicated a varying rate of exchange, suggesting that the participant was giving up lesser amounts of component B to obtain one unit of component A when advancing on the IC from top left to bottom right. For a more intuitive measure, we quantified the curvature by measuring the largest perpendicular distance between the IC and the line between the x-axis and y-axis intercepts (Figure 2E):

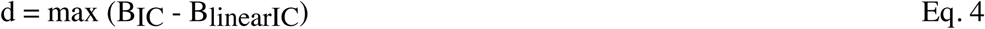

with d as maximal perpendicular distance (ml) (whereas β_3_ was a best-fitted, estimated parameter, and thus less conservative), B_IC_ as amount of component B on the IC (ml), and B_linearIC_ as amount of component B at the line connecting the x- and y-intercepts (constant amount of component A, x-axis; ml). This simplified curvature measure reflected the change in trade-off between the two components across the tested range of reward amounts, in ml of component B.

We used logistic regression on trial-by-trial choices to confirm that the measured choices were explained by bundle components rather than other factors. In a random-effects analysis, we fitted a logistic regression to the data from each individual participant and then averaged the obtained β coefficients and p-values across all participants. We used the following regression:

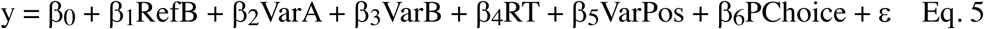

with y as either 0 or 1 (not-choice or choice of Variable Bundle), A and B as amounts of bundle components A and B (ml), RefB as amount of component B in the Reference Bundle (ml), VarA and VarB as amount of components A and B in the Variable Bundle (ml), RT as reaction time (ms; interval between appearance of the two bundle stimuli and participant’s computer button press), VarPos as left or right position of the Variable Bundle on the computer monitor relative to the Reference Bundle (0 or 1), and PChoice as choice in the previous trial (0 or 1) (Figure 2G). Each β coefficient was standardised by multiplication with standard deviation of the respective independent variable and division by standard deviation of y (dependent variable). A subsequent one-sample t-test against 0 served to assess the significance of the beta (β) coefficients in the population of the 24 participants.

Satiety may have occurred and could have affected the preferences for the two bundle components in an uncontrolled manner, even though the bundle rewards were only paid out on every fifth trial on average and were limited to a total of 200 ml. A prime suspected effect might have been differential devaluation between the two bundle components that would result in changed currency relationship between the two components. We assessed such change between the two components by searching for gradual change in instantaneous choice probability above and below the IPs over 6 repeated test steps of 2 × 12 trials each (4 IPs on each of 3 ICs; total of 144 choices; Figure 3). We calculated the instantaneous choice probability at each test step, separately above and below the IP, as:

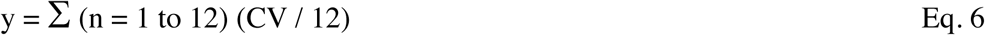

with y as instantaneous probability (*P =* 0.0 to 1.0), CV as choice of Variable Bundle (0 or 1).

To analyze the BDM bidding data, we first assessed the basic question whether the monetary bids increased for higher valued bundles (between revealed preference levels) but were similar for equally valued bundles (which constituted IPs) (along the same preference levels), using two-way Anova between and along preference levels, respectively, and confirmation by Spearman rank correlation analysis between preference levels. In addition, we performed a random-effects analysis with a general linear regression with a normal (Gaussian) link function on separate data from each participant and averaged the obtained β coefficients and their p-values across participants. We used the following regression:

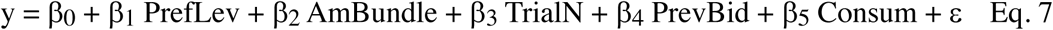

with y as monetary bid, PrefLev as revealed preference level (low, medium, high), AmBundle as summed ml amount of components A and B in the currency of component A (A + (k - (β_2_/β_1_)A + β_3_A) as in Eq. 3a), TrialN as trial number, PrevBid as BDM bid in previous trial (UK pence), and Consum as accumulated drinks consumption for component A and component B up to this point in the experiment (ml) (see Figure 4C). We included PrefLev and AmBundle as separate regressors to account for their distinction due to partial physical non-dominance of bundles on different preference levels, combined with currency differences between the two components. Each β coefficient was standardised by multiplication with standard deviation of the respective independent variable and division by standard deviation of y (dependent variable). A subsequent one-sample t-test against 0 assessed the significance of the beta (β) coefficients in all 24 participants.

Finally, we compared hyperbolically fitted BDM isolines directly with hyperbolically fitted revealed preference ICs (rather than with revealed preference levels just described), separately for each individual participant (note that the BDM data were acquired in the human imaging scanner and the IC data used in this analysis were acquired in a prior session with the binary choice task outside the scanner). This procedure required to present BDM bids on the same scale as revealed preference ICs. To this end, we fitted isolines of same BDM-bids in analogy to fitting same-preference ICs (see Figure 5A, B). We fitted a hyperbolic function to the measured mean BDM bids in analogy to Eq. 3:

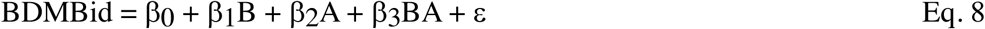

with β_1_ and β_2_ as regression slopes, and β_3_ as curvature coefficients, and A and B as amounts of components A and B (ml), respectively. Coefficients β_1_ and β_2_ were standardised by multiplication with standard deviation of components B and A, respectively (independent variables), and division by standard deviation of BDMBid (dependent variable). We obtained separate β coefficients from all participants and averaged them and their p-values across participants. A subsequent one-sample t-test used the individual beta (β) coefficients from all 24 participants to test overall significance against 0.

To compare BDM bids with ICs, we graphically displayed BDM isolines along which all mean BDM bids were equal. As a BDM isoline is a constant, we merged the constants offset (β_0_) and error (ε) into a common final constant k. To draw the BDM isolines, we computed the amount of component B as a function of component A from the derived equation:

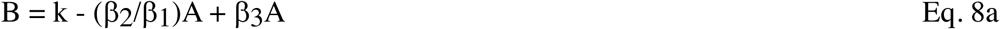

To display a three-dimensional map, we graphed colored BDM isoline zones on the z-axis as a function of the amounts of components A (x-axis) and B (y-axis) (see Figure 5A). For a two-dimensional map of BDM isolines, we plotted the preset amount of component A on the x-axis and the amount of component B computed from the isolines (Eq. 8a) on the y-axis (see Figure 5B). For comparison, we plotted the revealed-preference ICs on the same two-dimensional map using the same scale. We also compared numerically, separately for each participant, CIs and slope and curvature coefficients between hyperbolically fitted BDMBids (Eq. 8a) and hyperbolically fitted revealed preference ICs (Eq. 3a), using the paired Wilcoxon test.

### Comparison with monkeys

In order to compare the results of this human study across a closely related species, we re-analyzed existing data from a previous experiment on rhesus monkeys; all methods of this study have been described (Pastor-Bernier, Plott, & Schultz, 2017). That study tested in an analogous manner the same notions of revealed stochastic preferences stated above. Each monkey chose with a hand touching a horizontally mounted monitor between two visually presented bundles composed of the same two liquids (fruit juices or water) in varying amounts. We used analogous psychophysical procedures and statistics to estimate IPs and ICs (see Figure 6A-E). For the current comparisons, we assessed the accuracy of integration of the two option components by the monkeys with two measures (see Figure 6F-M): (1) the 95% CIs of the psychophysical fits to the choice probabilities used for estimating each IP, which indicated how well the animals had estimated the IPs (Figure 2A inset); (2) the CIs of the hyperbolic fits of ICs to all equally preferred IPs, which indicated the accuracy of the trade-off between the two bundle components that characterizes the value integration from both bundle components (Figures 2A, B; S1).

**Figure 6.**
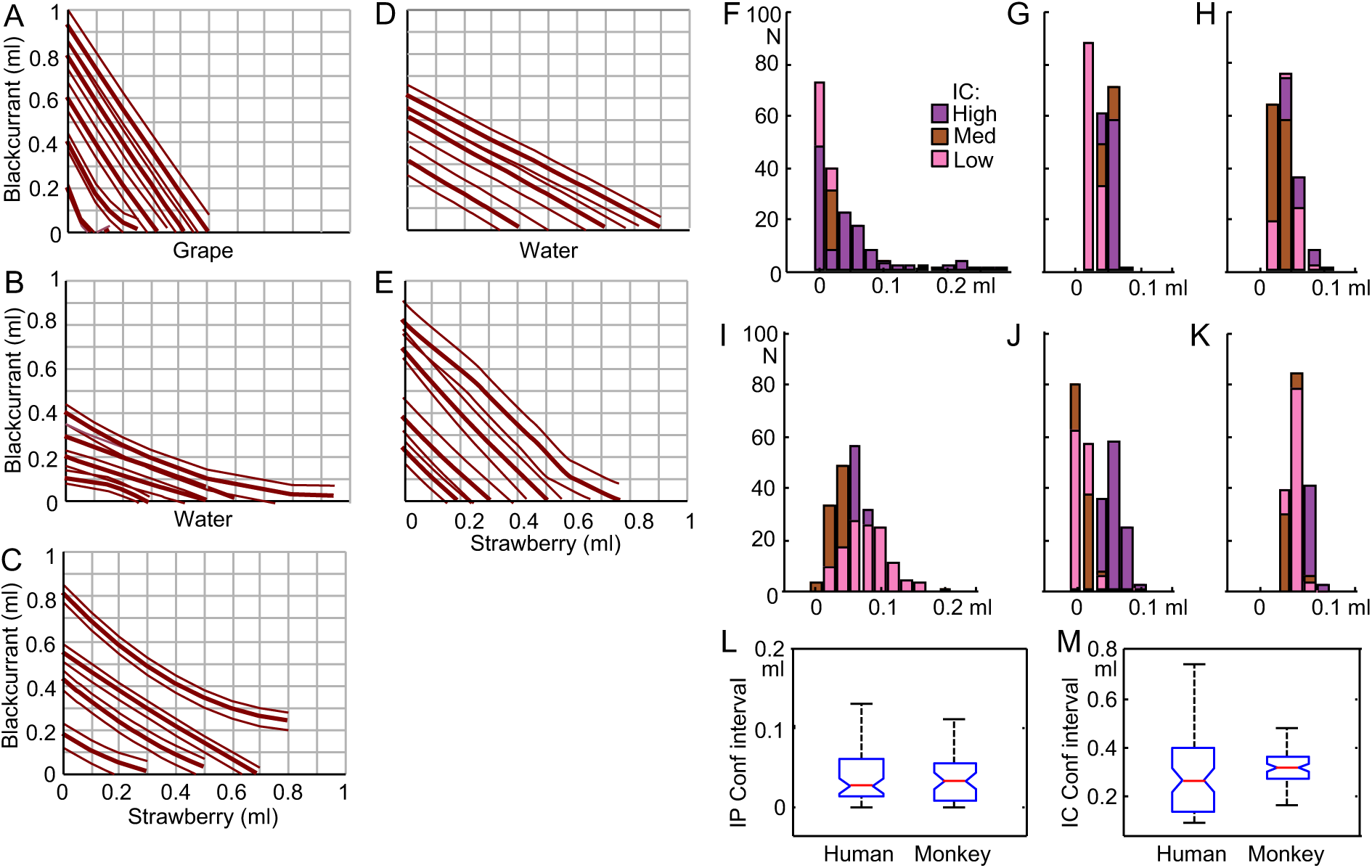
Comparison of choice accuracy with rhesus monkeys. (A - E) Empirically estimated indifference curves for five bundle types in monkeys. Heavy lines show best-fitting quadratic polynomials, thin lines show 95% confidence intervals of the least-mean-square fits to the averaged data. All data are from Pastor-Bernier, Plott, & Schultz, 2017. (A) bundle type (blackcurrant juice, grape juice) in monkey A (B) as A but for bundle (blackcurrant, water) (C) as A but for bundle (blackcurrant, strawberry juice) (D) as B but for monkey B (E) as D but for bundle (blackcurrant, strawberry) (F) Human participants: confidence intervals (CI, 95%) from psychophysical fits for estimation of 5 choice indifference points (IP) on each of 3 indifference curves (IC) (pink, brown, violet) from 24 participants (0.032 ± 0.005 ml; *N* = 360 IPs), normalized to reward range for monkeys (0 - 1.0 ml; milkshake). Inset applies to A - F. All columns start at bottom. (G - K) Monkey CIs (95%) for the five bundle types shown in A-E. *N* = 40 - 50 IPs / bundle type. To obtain same sample size as in humans, 360 randomly selected IPs were resampled 100 times and averaged in each of 360 bins, separately for each bundle type. (G) CIs (95%) for bundle (blackcurrant, grape) in monkey A (0.043 ± 0.008 ml of blackcurrant juice; mean ± SEM). (H) as G but for bundle (blackcurrant, water) (0.040 ± 0.006 ml). (I) as G but for bundle (blackcurrant, strawberry) (0.061 ± 0.011 ml). (J) as H but for monkey B (0.065 ± 0.016 ml). (K) as J but for bundle (blackcurrant, strawberry) (0.025 ± 0.013 ml). (L) CIs (95%) from psychophysical fitting for IP estimation. Box plots show medians (0.0310 ml milkshake normalized for humans; 0.0344 ml blackcurrant juice for monkey; *P* = 0.000461; CIs for *N* = 360 IPs each species; two-tailed Kolmogorov-Smirnov test; red), 25th and 75th percentiles (blue) and most extreme values (black). (M) CIs (95%) from hyperbolic fitting for IC estimation. Medians were 0.2658 ml milkshake normalized for humans; 0.3038 ml blackcurrant juice for monkey; *P* = 0.0018; CIs for *N* = 72 ICs each species (reduced for monkeys by random selection to correspond to 3 ICs in 24 humans); K-S test.

## Results

### Human revealed preferences follow the graphic scheme of indifference curves (IC)

We obtained individual ICs from each set of five equally revealed preferred IP bundles by hyperbolic fitting (Eqs. 3, 3a) (Figure 1E, F). Such an IC defined the trade-off between the two components of an equally preferred bundle: it indicated how much of component B a participant gave up for obtaining one more unit of component A without a change in utility. Thus, the IC characterised the orderly integration of both bundle components into a single-dimensional estimate. The continuous ICs were asymmetric between the x-axis and y-axis, indicating different subjective weighting of the two milkshakes; the convex IC curvature suggests that lower amounts of both milkshakes together were as much preferred as higher amounts of singular milkshakes (possibly reflecting gradually flattening, concave utility functions and/or complementarity between high-sugar and high-fat components). Although the ICs varied in slope and curvature between participants, the ICs of bundles with larger reward amounts were located farther away from the origin (for two example participants, see Figure 2A, B; for all participants, see Figure S1; the subjective, individual nature of ICs does not allow to assume a common preference and utility scale, which precludes averaging ICs across participants). The three ICs were well ordered and failed to overlap. Only the 95% confidence intervals (CI) overlapped partly in four of the 24 participants (17%).

Closer inspection of Figures 1F, 2A, B and S1 shows a typical feature of ICs. During the psychophysical assessment of individual IPs, the Variable Bundle was preferred to the Reference Bundle (choice *P >* 0.5) even when component B of the preferred bundle had lower amounts compared to the Variable Bundle at the y-axis anchor (Figure 1F, arrows), indicating partial physical non-dominance (notions 7 and 8; involving overcompensation by higher amount of component A). As with the trade-off, such preferences for bundles with one physically lower component indicate the integration of the values of both bundle components into single-dimensional utility.

Taken together, the maps of systematic and continuous ICs reflect well the integration of both bundle components into single-dimensional utility and preference relationships. As an alternative to these well integrated ICs, lexicographic preferences would not show such integration but follow only a single bundle component, manifested as lines parallel to one of the graph’s axes. Taken together, the true, multi-component-integrating ICs corresponded well to, and thus seemed to validate empirically, the intuitive graphic schemes of Revealed Preference Theory. The following tests will address the validations numerically.

### IC coefficients

The shape of ICs reflects the trade-off between the components and can be quantified by slope and curvature coefficients of hyperbolic fits (Eq. 3a). The global IC slope, between y-axis and x-axis intercepts, was measured as ratio of the two regression coefficients β_2_ / β_1_ in Eq. 3; it indicated how much the participant was globally willing to give up in order to obtain one unit of the other component; the measure reflected the relative utility (currency; global MRS) of the two bundle components. The IC slopes steeper than −45° indicated that a participant gave up a higher physical amount of component B (high-sugar, y-axis) for a smaller amount of component A (high-fat, x-axis), thus indicating higher subjective value of fatty than sugary milkshake per unit of ml. The IC slope was −71° ± 6.5° (mean ± standard error of the mean, SEM; range −45° to −76°; *N* = 24 participants; Figure 2C). The higher valuation of high-fat component A over high-sugar component B amounted to a factor of 3:1 in 18 of the 24 participants (75%). The predominantly asymmetric trade-off between the two milkshake components documents that each component contributed to bundle preference in its own distinct way.

The IC curvature showed substantial convexity in 18 of the 24 participants (75%), as indicated by β_3_ coefficients from the hyperbolic fit (Eq. 3) that were significantly larger than 0.0 (8.89; mean ± 5.9 SEM; *p<*0.05, t-test; Figure 2D). For graphically assessing IC curvature, we measured the distance between the IC center and a straight line connecting equally revealed preferred bundles at the x-and y-intercepts, in units of ml on the y-axis (Figure 2E). This distance ranged from 0.09 ml (quasi-linear IC) to 3.76 ml (most convex IC) (mean of 1.28 ml ± 0.19 SEM; Figure 2F). The distribution of the IC distance was overall similar to that of the β_3_ curvature coefficient from Eq. 3. The two highest histogram bars in Figure 2F show data from six participants with rather similar, considerably convex IC, and from six other participants with rather similar but quasi-linear IC. Thus, the coefficients confirmed numerically the well-ordered nature of the ICs representing revealed preferences.

### Control for other choice variables

To test whether the choices reflected the components of the bundles rather than other, unrelated factors, we performed a logistic regression analysis separately for each of the 24 participants, using the following regressors: amount of each bundle component, reaction time, Reference Bundle position on participant’s monitor, and previous trial choice (Eq. 5). Median reaction times were 2.10 s and 1.98 s with > 1 ml milkshake difference above or below IP, respectively, and 2.58 s with < 1 ml difference from IP. The standardised beta (β) coefficients and p-values were assessed for each participant and then averaged across all 24 participants; they demonstrated that the choice of the Variable Bundle was negatively correlated with the amount of component B in the Reference Bundle (RefB: *β* = −0.43 ± 0.16, mean ± SEM; *P =* 0.020 ± 0.005) (component A was constant 0.0 ml, see Methods) and positively correlated with both components A and B in the Variable Bundle (VarA: *β* = 0.67 ± 0.16; *P =* 0.009 ± 0.004; VarB: *β* = 0.94 ± 0.33; *P =* 0.012 ± 0.009) (Figure 2G). The beta (β) coefficients for these three variables differed significantly from 0 (*P =* 0.012, *P =* 0.00088 and *P =* 0.00028, respectively; one-sample t-test), confirming the validity of the β’s. Thus, the Variable Bundle was preferred with increasing amounts of either of its components and with lower amounts of the Reference Bundle. The result suggests that both bundle components, rather than a single component alone, were important for the observed preferences. The remaining variables, including reaction time, position of Reference Bundle on the monitor and previous trial choice, failed to significantly account for current choice of the Variable Bundle (*P =* 0.754 ± 0.003 to *P =* 0.988 ± 0.290). Thus, the revealed preference relationships concerned the bundles with their two components rather than other task factors.

To assess potential consumption effects derived from the flow of the milkshake during the experiment, we searched for signs of satiety. We followed choice probability across the total test duration in each of the 24 participants. We selected two bundles that were situated above and below the IP, respectively. These two bundles contained high-sugar low-fat (above IP) and low-sugar high-fat (below IP) milkshakes. We plotted choice probability over six repeated test steps. Choice probabilities fluctuated without conspicuous upward or downward trend (Figure 3) and varied only insignificantly (above IP: F (5, 41) = 0.28, *P* > 0.05; below IP: F (5, 41) = 1.53, *P* > 0.05; one-way repeated measures Anova). Even at the final, sixth step, choice probability differed only insignificantly from any other step. Thus, the revealed references did not seem to be importantly confounded by satiety for neither sugar nor fat within the amounts and concentrations used in our experiment.

### Internal validation of IPs and Ics

We assessed the contribution of individual IPs to the hyperbolically fitted ICs with three tests. (1) Using a leave-one-out analysis, we compared ICs fitted to all five IPs with ICs fitted with one IP left out and found good correspondence in all of four tests (Figure S2). (2) Using a previously developed single-dimensional linear support vector machine (SVM) algorithm (Grabenhorst, Hernadi, & Schultz, 2016; Tsutsui, Grabenhorst, Kobayashi, & Schultz, 2016) and (3) a two-dimensional linear discriminant analysis (LDA), we assessed the accuracy of reversely assigning individual IPs to their original preference level. Both decoders reported between-IC accuracies largely in the 70% - 100% range, and the LDA showed only random distinction along-ICs (Figure S3; Table S3 left); as a control, shuffled data did not discriminate between preference levels (SVM accuracies of 44.7% - 54.6%; Table S4 left). These procedures confirmed that the hyperbolically fitted ICs captured the IPs consistently and provided valid representations of the revealed preferences for two-component choice options. For details, see Supplementary Material.

### Mechanism-independent validation

The ordered representation of revealed preferences can be further validated by comparison with utility inferred from a different estimation mechanism. We used a monetary Becker-DeGroot-Marschak (BDM) auction-like bidding task that estimates participants’ utility on a trial-by-trial basis. The property of truthful revelation (incentive compatibility) makes BDM an indispensable tool for experimental economics and explains its increasing popularity in neuroscientific studies of human decision making (Plassmann, O’Doherty, & Rangel, 2007; Medic et al., 2014). We ran this task independently from the binary choice task in an fMRI scanner (fMRI data to be reported separately) and obtained BDM bids in UK pence for each of the 15 IP bundles (Figure 4A). Although BDM bids are assessed as single shots, we repeated each trial 12 times to approach the nature of stochastic choice for IP estimation. We compared the 12-trial averaged bids to preference levels rather than to ICs to avoid fitting inaccuracies.

A two-way Anova showed that BDM bids varied between the three revealed preference levels in all 24 participants (*P* = 5.37 × 10^−40^ to *P* = 5.38 × 10^−116^) but rarely between the five equally revealed preferred bundles (IPs) on each level (*P* < 0.05; except for three participants) (Figure 4B; red, green, blue; Table S5). This variation pattern was confirmed with separate one-way Anovas between the three preference levels (*P* = 0.038534 to *P* = 6.86 × 10^−16^) and the five IPs along the same preference levels (*P* = 0.055 to *P* = 0.951; except for three participants *P* < 0.05). A Spearman rank analysis confirmed and refined the result between preference levels; it showed a positive monotonic correlation between bids for bundles on the same preference level (means from all five bundles) and the three levels (rho = 0.60 ± 0.05; mean ± SEM; *N* = 24 participants; *P <* 0.01). Thus, the Anovas and Spearman correlation showed showed higher BDM bids between the three increasing revealed preference levels but mostly similar bids for bundles on same preference levels (with the Anova being sensitive to bundle amounts, as shown by increasing bids between levels). This BDM result confirmed the pattern of revealed preference seen with binary choice: higher value for bundles on higher preference levels, and similar value at trade-off between bundle components.

A random-effects analysis separately for each of the 24 participants (Eq. 7) confirmed the relationship of BDM bids to preference level (PrefLev: *β* = 0.47 ± 0.09, mean across all 24 participants ± SEM; *P =* 0.016 ± 0.015; β-coefficient difference from 0: *P =* 0.000026, one-sample t-test) and bundle amount (AmBundle: *β* = 0.15 ± 0.13; *P =* 0.020 ± 0.017; *P =* 0.0278; AmBundle varied separately from PrefLev due to partial physical non-dominance of bundles on different preference levels combined with currency differences between components), rather than trial number (TrialN: *β* = −0.10 ± 0.25; *P =* 0.726 ± 0.354), previous trial bid (PrevBid: *β* = 0.12 ± 0.11; *P =* 0.676 ± 0.427) or consumption history (Consum: *β* = 0.12 ± 0.11; *P =* 0.224 ± 0.185) (all β’s were standardised) (Figure 4C). A specific analysis (Eq. 8) demonstrated that the BDM bids reflected both bundle components (component A: *β* = 0.6534 ± 0.0866, mean ± SEM; *P =* 0.0324 ± 0.0150; β difference from 0: *P =* 1.1613 0884 × 10^−7^, one-sample t-test; component B: *β* = 0.6425 ± 0.0585, *P =* 0.0289 ± 0.0202; β difference from 0: *P =* 1.2770 × 10^−10^). Thus, the BDM bids followed well the revealed preference levels and took both bundle components into account.

Using SVM and LDA decoders, we assessed the accuracy of reversely distinguishing original preference levels from individual BDM bids. Both decoders reported discrimination accuracies largely in the 50% - 70% and the 88% - 100% range, respectively, between preference levels (Table S3 right), but the LDA showed only random distinction between bundles on same preference levels (43-51%) (Figure S4); shuffled data did not discriminate between preference levels (SVM accuracies of 45.8% - 54.7%; Table S4 right). Thus, decoding accuracy confirmed well the two-dimensional scheme of revealed preference represented by the ICs. For details, see Supplementary Material.

A stronger mechanism-independent validation of the IC scheme may be achieved by direct graphic comparison between BDM bids and hyperbolically fitted ICs. To this end, we estimated isolines that were fit to BDM bids using Eqs. 8 and 8a and compared them with ICs that had been hyperbolically fitted to IPs (Eqs. 3, 3a). The BDM isolines represented BDM bids from 12 averages, and the revealed preference ICs derived from fits to IPs estimated from 42 stochastic choices. The BDM isolines increased for IPs on increasing preference levels (farther away from the origin), but were similar for IPs on the same revealed preference level (Figure 5A). The BDM isolines matched the revealed preference ICs within their 95% CIs in every one of the 24 participants (Figure 5B). Statistical comparisons showed significantly higher CIs of BDM isolines compared to revealed preference ICs (Figure 5C; *P <* 0.0884 × 10^−8^, Wilcoxon paired test; *N* = 24 participants). Despite their larger variability, both BDM isoline slope and curvature coefficients, derived from the respective β_2_ / β_1_-ratio and β_3_ regression coefficient in Eq. 8, failed to differ significantly from the respective slope and curvature coefficients of revealed preference ICs (Eq. 3; Figure 5D, E; both *P >* 0.05, Wilcoxon test on BDM vs. IC coefficients paired from each participant; *N* = 24). Thus, despite larger variability, the BDM bids matched well the revealed preference ICs when assessed in a comparable way.

Taken together, the mechanism-independent validation with BDM bids provided good confirmation of the graphic representation of revealed preferences by the ICs and thus strengthens the validity of these conceptual schemes.

### Performance comparison with non-human primates

The systematic IPs reflecting the trade-off between the two bundle components resulted in well-ordered ICs of our human participants that reflected systematic integration of utilities of all option components. The question arises to what extent these basic measures of multi-component integration in humans compared with non-human primates whose lack of verbal interaction makes their behavior primarily dependent on experienced reinforcement. Using the same economic concepts and experimental design as in our human participants, we had tested two rhesus monkeys during several months and estimated > 600 IPs that conformed to three to six convex or linear, negatively sloped ICs with each of five different types of two-component bundles containing blackcurrant juice, grape juice, water and strawberry juice (Figure 6A-E; Pastor-Bernier, Plott, & Schultz, 2017). The ICs were consistent, as shown by out-of-sample prediction, transitivity and independence of number of choice options, thus reflecting the animals’ extended laboratory experience and allowing comparison with human performance.

We used two measures for the decision process underlying revealed preferences for multi-component choice options. First, to assess the accuracy of the estimated IPs, we computed the 95% CIs of the psychophysical fits to the choice probabilities when assessing individual IPs (which were used for constructing ICs). The CIs for IPs in humans were in a similar range but slightly smaller in comparison with all but one of the five bundle types in monkeys (Figure 6: compare F with G-K); the higher human accuracy was overall significant (*P* = 0.000461; Kolmogorov-Smirnov test; Figure 6L). Second, we tested the accuracy of the fit of ICs to the IPs that were derived from the trade-off between the two bundle components; this trade-off critically reflected the integration of the two bundle components into single-dimensional measures. To this end, we computed the CIs of the hyperbolic IC fits to the estimated IPs. A narrow CI would indicate a good match of ICs to IPs and thus a good trade-off and utility integration between the two bundle components. The human participants showed slightly smaller CIs, and thus higher accuracy, than the monkeys (*P* = 0.0018; Figure 6M).

Thus, despite differences in number of participants (*N* = 24 for humans, *N* = 2 for monkeys), task experience (single days for humans, several months for laboratory monkeys) and reward (milkshakes for humans, fruit juices and water for monkeys), the CI comparisons for IPs and ICs suggested comparable, although slightly better, choice accuracy in humans compared to the monkeys. Besides these encouraging results, the species comparison suggests robust correspondence of empirical ICs to the schematic graphs, thus validating the theoretical concepts.

## Discussion

This study used formalisms of Revealed Preference Theory to empirically investigate decisions for multi-component rewards. We measured stochastic choices of small, tangible and immediately consumable milkshake rewards that were delivered in repeated trials in a controlled, non-verbal laboratory setting. The estimated ICs were systematic and corresponded well to graphic economic schemes that are used for interpreting human choices (Laidler & Estrin, 1989; Kreps, 1990; Varian 1992; Mas-Colell, Whinston & Green, 1995; Perloff, 2009). The similarity in performance accuracy with monkeys suggested maintained preference mechanisms across evolutionarily gaps. Further comparisons between the two species, in particular concerning the nature of preferences indicated by IC shape, should be done with caution, as IC shape was also influenced by bundle composition that differed between the species. The experimental feasibility opens the possibility for concept-driven, empirical neuroimaging studies of rational and irrational behavior in normal and brain-damaged individuals.

Our experimental controls and comparisons attested to the validity of these measurements; the ICs were asymmetric and nonlinear, failed to overlap, complied with decoder predictions and correlated with independent BDM bids. The integration of multiple bundle components was particularly evident in the continuous, graded trade-off along ICs. Preference was unchanged when reduction in one bundle component was compensated by increase in another bundle component, and bundles with one smaller component could even be preferred to bundles with a larger component when the other component compensated (partial physical non-dominance; Figure 1F). Such choices suggested proportionate contributions of individual bundle components.

The representation of vectorial, multi-component choice options by single-dimensional neural signals is an open issue in experimental neuroeconomics. Our behavioral tests provide a formal, concept-driven foundation for investigating such signals in humans. The immediately consumed, small payouts facilitate comparisons with animal studies and control for temporal discounting. The tangible payouts after small sets of trials do not rely on language and should reduce mental ruminations about future rewards and assure reliable cooperation by the participants, thus reducing interfering and confounding brain activity. Future neuroeconomic work on underlying decision mechanisms should particularly benefit from the empirical estimation of whole maps of well-ordered ICs derived from multiple IPs that conform to predictive mathematical functions, thus avoiding to test preferences for a few bundles with limited general validity. For example, different neurons in the orbitofrontal cortex of monkeys combine both bundle components into a common scalar neural signal or code each component separately (requiring later integration for contribution to the decision; Pastor-Bernier, Stasiak, & Schultz, 2019). The systematic ICs may also help to investigate neural underpinnings of specific theories, such as the switching of attentional processes between components conceptualised by multialternative decision field theory (Roe, Busemeyer, & Townsend, 2001). Finally, our detailed ICs would be helpful for investigating neural mechanisms underlying inconsistent decision making when choice options are added (independence of irrelevant alternatives, IIA), beyond the decoy effects already being addressed (Chung et al., 2017; Gluth, Hotaling, & Rieskamp, 2017).

Our multi-trial approach corresponds to the standard psychophysical elicitation of choice functions (Green & Swets, 1966; Sutton & Barto, 1998), allows comparison with animal studies (Kagel et al., 1975; Pastor-Bernier, Plott, & Schultz, 2017) and was tailored to the statistical requirements of neural studies. In humans and monkeys, these methods deliver systematically varying, graded choice probabilities rather than single, all-or-none selection. The visible trial-by-trial variations are assumed to reflect underlying random processes that make the choice process stochastic, as captured by well worked-out stochastic choice theories that facilitate data interpretation (Luce, 1959; McFadden & Richter, 1990; McFadden, 2004; Stott, 2006). We appreciate that our multi-trial schedule is at odds with the frequently employed, standard, single-shot, deterministic assessment of ICs in experimental economics (Thurstone, 1931; MacCrimmon & Toda, 1969; but see Mosteller & Nogee, 1951), and the obtained consistent and robust ICs seem to validate the approach.

Economic choice often involves substantial but imaginary sums of money or consumer items, or random, singular payouts (Simonson, 1989; Tversky & Simonson, 1993; Rieskamp, Busemeyer, & Mellers, 2006). By contrast, our payout schedule was tailored to requirements of neural studies in humans and animals and allowed immediate reward consumption over many trials (while controlling for satiety). These behavioral choices resembled small daily activities, such as consuming snacks and drinks, and were met by the good motivation of our participants. In this way, we obtained well-ordered ICs without requiring hypothetical items or large sums of imagined money.

Previous investigations of multi-component economic choice have revealed several inconsistencies, including reference biases, difference between willingness-to-pay and willingness-to-sell, and violation of independence of irrelevant alternatives such as similarity, compromise, asymmetric dominance and attraction effects (Tversky, 1972; Knetsch & Sinden, 1984; Simonson, 1989; Tversky & Simonson, 1993; Bateman, I., Munro, Rhodes, Starmer, & Sugden, 1997; Rieskamp, Busemeyer, & Mellers, 2006). These phenomena may be due to revealed preferences being constructed at the time of choice rather being fixed (Payne, Bettman, & Schkade, 1999; Simonson, 2008; Dhar & Novemsky, 2008; Kivetz, Netzer & Schrift, 2008; Warren, McGraw, & Van Boven, 2011) or reflect the adaptive nature of biological processes (Soltani, De Martino, & Camerer, 2012; Li, Michael, Balaguer, Castanon, & Summerfield, 2018). We aimed to avoid interference from adaptive processes by designing stable and highly reproducible test conditions in a well-controlled laboratory environment, non-reliance on verbal report, single, uninterrupted test sessions, singular changes of bundle components, constant direction of testing (from top left to bottom right on the revealed preference map) and preventing satiety by limiting total milkshake intake to 200 ml. These measures may explain why our IPs remained stable over successive testing steps. We used the exact same conditions for eliciting BDM bids, which may have facilitated their correspondence to revealed preference ICs. With these testing conditions, we avoided known compromising factors that might hinder identification of basic factors underlying choice irregularities.

Our experiment used a limited range of closely related, basically substitutable reward types; thus, the validity of our results is restricted to that range and might not necessarily generalize to bundles with more varied kinds of rewards. Apes, monkeys, dogs and pigeons may display seemingly irrational preference patterns for bundles of food items with very different values. According to the ‘selective value effect’, an animals may assign value primarily or exclusively to a preferred reward item and partly or completely forego less desired items, rather than revealing more graded preferences (Silberberg, Widholm, Bresler, Fujita & Anderson, 1998; Pattison & Zentall, 2014). One possible explanation might be that consumption of a less preferred reward item would delay acquisition of the next preferred reward item (Beran, Ratliff, & Evans, 2009), but many other reasons might exist (Zentall, 2019). However, none of these tests assessed ICs, and the results are therefore difficult to compare with ours. With more formal testing, some of these anomalies may have been comparable to lexicographic preferences that concern only a single component and become evident by ICs that run parallel to one of the axes; our study failed to find lexicographic preferences. Further, our study only observed preference ratios up to 3:1, which suggested reasonable acceptance of both bundle components rather than generating selective value effects (see the IC slopes in Figures 2A, B; S1). Our parallel study on monkeys tested a wider range of liquid rewards and found similar two-dimensional ICs and no evidence for major violation, refuting possible reasons in species difference. Taken together, choices of multi-component bundles may be more prone to irregularities than choices of single-component options, but our experiment was too limited to address this tissue in an exhaustive manner.

The coefficients of hyperbolic fits to the ICs characterize numerically the representation of revealed preferences. The slope coefficient indicates the relative weighting of the two bundle components. For example, the amount of equally revealed preferred single milkshakes (graphed at the respective x-axis and y-axis intercepts of the two-dimensional map) was lower for high-fat (component A) than high-sugar (component B) milkshakes in our participants, which was represented by a IC slope steeper than −45 degrees; thus, participants would have revealed preferred high-fat over high-sugar milkshakes if they came in same amounts. Such asymmetric IC slopes are also seen with various bundles in monkeys (Pastor-Bernier, Plott, & Schultz, 2017). Key reasons for the visible asymmetry on the graphic axes may be the physical amount scale and the absence of a simple common physical scale for the fat and sugar content of the milkshakes. Further, the slope for identical bundle compositions varied between our participants, which demonstrates an additional subjective component in revealed preferences. Despite these scaling and subjectivity issues, our estimated ICs had well-ordered slopes and failed to overlap.

The convex IC curvature in most participants indicated that disproportionately smaller amounts of combined fat-sugar milkshakes (at IC center) were equally revealed preferred as larger amounts of milkshakes containing primarily fat or sugar (closer to the axes). The BDM isolines confirmed the convex curvature for probably the same reasons. A previous study on food snacks also found higher BDM bids for fat-sugar combinations than for fat or sugar alone, despite similar calories (DiFeliceantonio et al., 2018). However, not all ICs need to be convex; bundles with one unattractive component showed concave ICs in primates (Pastor-Bernier, Plott, & Schultz, 2017). The IC convexity may be ascribed to concave utility of each bundle component (Perloff, 2009); more of the same component (closer to IC x- or y-intercepts) has lower marginal gain, and therefore the decision-maker is willing to trade in more of an abundant, undervalued component for one unit of a less abundant, more valued component. Thus, the defining slope and curvature coefficients suggest that the ICs reliably represented well-ordered preferences for the milkshake bundles.

The estimating mechanism of BDM bids differed substantially from that underlying revealed preference ICs. Bidding occurred along a single scale and involved the physical movement of a cursor rather than hitting choice buttons between two simultaneous options. BDM bidding is incentive compatible, such that erroneous or deceiving bids lead to suboptimal outcome, as conceptualised by the expected cost of misbehavior (Lusk, Alexander, & Rousu, 2007); bidders should state exactly their subjective value to avoid paying an exaggerated price (with too high bids) or foregoing the desired object (with too low bids). With these properties, BDM bidding constituted a well-controlled, authoritative test for eliciting true subjective values and provided a useful validation mechanism for preferences revealed by binary choice. Indeed, the obtained SVM- and LDA-consistent BDM bids followed the preference scheme of ICs, namely higher bids for revealed preferred bundles and similar bids with choice indifference despite varying bundle composition. Most strikingly, hyperbolically fitted BDM isolines closely resembled the hyperbolically fitted revealed preference ICs in graphics and numerics (Figure 5). The only notable difference was higher BDM bid variability. Our data align well with, and extend, the previously noted correspondence between binary choices and BDM bids for single-component options and bundle choices on paper or via verbal communication (Roosen, Marette, & Blanchemanche, 2017). This correspondence is particular interesting in light of conflicting and unclear accounts of economic choice and utility, such as the unresolved distinction between inherent and constructed revealed preferences (Payne, Bettman, & Schkade, 1999; Simonson, 2008; Warren, McGraw & Van Boven, 2011) and the fundamental question whether utility, supposedly underlying both revealed preferences and BDM, may not be a required inferred variable, and choices may simply involve heuristics (Vlaev, Chater, Stewart, & Brown, 2011; Piantodosis & Hayden, 2015). Whatever the answer might be, the BDM bids validated the empirical assessments of the ICs and confirmed their representation of revealed preferences.

The revealed preferences in humans indicated a similar level of integration of the two-component choice options as in rhesus monkeys (Pastor-Bernier, Plott, & Schultz, 2017) and, in a more general way, also observed in rodents (Kagel et al., 1975; Kagel, Battalio, & Green, 1995). In two monkeys, the estimated IPs conformed to three to six convex or linear, negatively sloped ICs with five types of two-liquid bundles. Preferences for other bundles were characterized by concave or positively sloped ICs reflecting satiety or disfavored juices. The validity of preference representation by the consistent and orderly ICs in monkeys was shown by out-of-sample prediction from homothetic polynomials, various transitivity tests, axiomatic compliance with independence of number of choice options, and classifier prediction. In contrast to the unexperienced humans, the monkeys had several months of training in well-controlled, stable laboratory conditions. The reward difference (milkshakes in humans vs. juices in monkeys) and the difference in IC estimation (unidirectional IP progression vs. random alternation) failed to prevent the general similarity of ICs between the two species, despite known hysteresis issues (Knetsch, 1989). Only quantitative differences were observed, with higher precision in humans, and it remains to be seen which of the different factors might account for these differences: cognition, task experience and/or other factors. Thus, the robustness of performance across species is reassuring for the validity of our experimental procedures and demonstrates the basic and precise nature of this economic decision process with revealed stochastic preferences for multi-component choice options.

## Acknowledgements

We thank Charles R. Plott for discussions and conceptual support, Steve Edgley for help and logistic support, Simone Ferrari-Toniolo for comments on experimental economics and Francine Ganter-Restrepo for engineering expertise. This work was supported by the Wellcome Trust (WT 095495, WT 204811).

## SUPPLEMENTARY MATERIAL

### Supplementary Methods

#### Leave-one-out analysis of Ips

We used a leave-one-out analysis to assess the meaningful representation of revealed preferences by the fitted ICs by testing the accuracy of the hyperbolic IC fit to the IPs. In this analysis, we removed one IP per IC (but not the initial Reference Bundle set at x = 0) and fitted an IC again with the same hyperbolic model as for the main IC fitting (see Methods, Eqs. 3, 3a). In total, for each original IC, we fitted 4 new ICs, each one leaving out a different IP. For each new IC, we assessed the deviation between the left-out IP and the refitted IC. We measured this deviation as difference of component B between the y-axis position of the original (but now left out) IP and the y-axis position of the refitted IC with the IP left out, at the same x-position (Figure S2B):

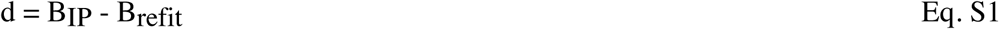

with d as difference (in ml; y-axis), B_IP_ as amount of component B of the left-out IP (ml), and B_refit_ as amount of component B on the refitted IC (ml). Thus, a difference of 0 ml suggested that removal of one IP did not affect the shape of the IC at all, thus indicating an excellent representation of revealed preferences by the fitted ICs, whereas any difference unequal to 0 ml quantified the degree of inaccuracy of this representation.

#### Decoder analysis of preference levels

To confirm the contribution of each IP to the two-dimensional representation of revealed preferences, we determined the accuracy (in percent correct) with which a randomly selected bundle, defined by the amounts of the two components A and B (in ml), could be assigned to its original revealed preference level as opposed to any one other level (binary distinction). By definition, each bundle that was psychophysically estimated to be as much revealed preferred as the Reference bundle constituted an IP; all bundles to which participants were choice indifferent against the same Reference Bundle constituted a series of IPs. In our experiment, three different Reference Bundles defined three preference levels (low, medium, high: component B: 2.0 ml, 5.0 ml or 8.0 ml, respectively; component A was always 0.0 ml; Figure S3A). The decoder used as inputs only bundles at the psychophysically estimated IPs (to which an IC was fitted using Eqs. 3, 3a), rather than bundles positioned on the fitted ICs.

Our main test employed a binary support vector machine (SVM) decoder separately on each individual participant. We used similar methods as previously described for predicting choice from neuronal activity (Tsutsui, Grabenhorst, Kobayashi, & Schultz, 2016). The SVM algorithm considered 5 IP bundles from each of 2 revealed preference levels (total of 10 IPs that had been assessed 12 times at each position in each participant) (Figure S3A). Each of the 2 preference levels was associated with a matrix of 2 columns (containing the x- and y-coordinates of bundle components A and B, respectively) and 5 rows (containing the 5 bundles). The 5 bundles were randomly selected (with replacement) from 60 bundles on each level (due to the random procedure with replacement, some bundles may have entered the algorithm multiple times, and not all five bundles may have been used for a given analysis). We left out 1 randomly selected bundle from the 10 bundles, trained the SVM algorithm with the remaining 9 bundles, and assessed whether the SVM decoder assigned the left-out bundle to its original revealed preference level or to another level. Thus we used 90% of the data for training the decoder and 10% for testing its classification performance. We repeated this procedure 10 times with the same selected 2 × 5 bundles but with a new randomly selected left-out bundle and calculated decoder accuracy as percent correct classification in these 10 trials. We repeated the random selection of the 2 × 5 bundles and the 10-trial accuracy assessment 150 times. For final decoding accuracy, we averaged the percentages from these 150 iterations (Table S3 left). We applied this procedure separately to all three possible combinations of two revealed preference levels (i. e. low and medium, medium and high, low and high). For assessing chance decoding, we shuffled the 2 × 5 matrix. Our earlier work has shown that increasing the number of analysis trials from 10 to 20 resulted in similar accuracy (Grabenhorst, Hernadi, & Schultz, 2016; Tsutsui, Grabenhorst, Kobayashi, & Schultz, 2016). The SVM was implemented with custom written software in Matlab R2015b (Mathworks) using the functions *svmtrain* and *svmclassify* with linear kernel (our previous work had shown that use of nonlinear kernels did not improve decoder performance; Tsutsui, Grabenhorst, Kobayashi, & Schultz, 2016).

We supplemented the SVM procedure with binary linear discriminant analysis (LDA) that provided visualization of the different levels of revealed preference (Figure S3). We used the same IPs and the same data matrices as for the SVM analysis (and the same IPs as used for the hyperbolic fitting of the three indifference curves, ICs). We obtained two variances; the discriminant 1 eigenvector captured the best separation between the three revealed preference levels as ‘between-level variance’ (colors in Figure S3); the discriminant 2 eigenvector captured the best within-level separation between five bundles on each of the three preference levels as ‘within-level variance’ (symbols in Figure S3). The results indicate visually the discrimination accuracy on the two axes of the two-dimensional plots. We also assessed the numeric accuracy of decoding as percent of correctly assigning a randomly selected bundle to its original revealed preference level. The decoder used the Matlab functions *fitcdiscr* and *predic* on z-normalised data from individual participants. For the LDA, our limited data required pooling from multiple participants. As revealed preferences are private and subjective, and therefore difficult to compare between individual participants, the LDA results should be considered as merely supportive and not as stand-alone data.

#### Decoder analysis of BDM bids

To assess the internal consistency of BDM bids, we used binary SVM analysis on bids from individual participants in analogy to SVM decoding of bundles according to preference levels. We tested the same IPs as used for hyperbolic IC fitting (Figure 4A). Each of the 2 preference levels was associated with a matrix of 1 column (containing the bids to each bundle) and 5 rows (containing the 5 bundles). The remainder of the bundle selection, repetition procedure and data shuffling was identical to that used for the SVM decoding for preference levels (see above). Thus, the SVM decoder for BDM bids assessed the accuracy with which the left-out bundle belonged to its original revealed preference level. We supplemented the SVM analysis of the BDM bids with analogous LDA for supportive visualization.

### Supplementary Results

#### Leave-one-out analysis of Ips

We performed a leave-one-out analysis to assess the contribution of individual IPs to the ICs obtained from hyperbolic fits to the empirically estimated IPs in humans. We removed one IP at a time from the set of five IPs per IC (except the initial Reference Bundle at x = 0), and then refitted each IC with the remaining four IPs using the hyperbolic model, separately for each IC and each participant (total of 3 ICs x 24 participants = 72 ICs, with 4 IPs x 72 ICs = 288 IPs; see Supplementary Methods). We found consistency in the refitted ICs in four measures (Figure S2). First, none of the 72 refitted ICs overlapped with the refitted ICs at different levels in the same participant, thus demonstrating maintained IC separation despite one left-out IP. Second, none of the 72 refitted ICs overlapped with the 95% CIs of original ICs at different levels, confirming IC separation despite one left-out IP. Third, most refitted ICs (66 of 72 ICs, 92%) fell inside the 95% CIs of the original ICs, and the remainder curves (6 of 72 ICs, 8%) showed only some portions outside the 95% CIs of the original ICs, thus refuting possible overweighted influence of individual IPs on ICs. Fourth, numeric comparisons showed only insignificant deviations between refitted ICs and the IPs that had been left out when refitting the curves (vertical distance of 0.05 ± 0.13 ml in all 24 participants; mean ± standard error of the mean, SEM; *N* = 336; *P =* 0.98 against normal distribution; t-test), confirming absence of overweighted IP influence on ICs. These four results suggest that the hyperbolically fitted ICs captured the IPs consistently and provided valid representations of the revealed preferences.

#### Decoder analysis of preference levels

We used a single-dimensional linear support vector machine (SVM) decoder as different statistical procedure to confirm the contribution of each IP to the two-dimensional representation of revealed preferences by ICs. In each participant, we set a given test bundle to one of the psychophysically determined IPs (Figure S3A) and assessed the accuracy with which the decoder assigned that bundle to its original preference level (each preference level was defined by a series of empirically estimated IPs but was not a fitted IC). SVM decoding accuracy ranged largely from 70% to 100% (*P* = 2.055 × 10^−101^), although a few lower values were observed (Table S3 left); shuffled data failed to discriminate between preference levels (accuracies of 44.7% - 54.6%; Table S4 left).

We supplemented the SVM analysis by visualization of decoding using two-dimensional linear discriminant analysis (LDA). The considerable amount of data necessary for reasonable LDA required us to pool data from several participants, which violates a basic tenet of economic theory that prohibits pooling of subjective preferences across individual participants. To somewhat contain expected inaccuracies, we normalised IPs across participants (z-score normalization for reward B along the y-axis; reward A had been experimentally set to identical values on the x-axis) and restricted the analysis to specific subsets of participants. The LDA confirmed the SVM results in all participant subsets (Figure S3); the first linear discriminant assigned bundles to the three revealed preference levels, as shown by spatial separation of the three colored groups, with a numeric accuracy of 80-100% (*P* = 1.148 × 10^−97^). By contrast, the second discriminant failed to accurately assign bundles to different positions on same preference levels, as shown by the mix of the five shapes representing bundle position. These characteristics were seen in six participants whose fitted ICs showed the highest similarity in convexity (Figure S3B, C), in six participants with linear ICs (Figure S3D, E) and, for comparison, in all 24 participants (Figure S3F, G) (for distinction of participants based on IC curvature, see two highest bars in Figure 2F). Thus, the two-dimensional LDA decoding followed the fundamentals of ICs: preference for bundles on higher ICs but indifference along ICs.

Taken together, the two decoders confirmed three distinguishable levels of IPs, and LDA in addition confirmed indifference between IPs on same levels. As these IPs constituted the basis for hyperbolic fitting of the three ICs, the decoder results validated also the fitting procedure and confirmed the representation of revealed preferences by the empirically estimated ICs.

#### Decoder analysis of BDM bids

We used decoders to test the distinction of bundle position between but not along preference levels. Using BDM bids, a binary SVM decoder showed good accuracy of assigning a test bundle to its original preference level in individual participants (mostly 50-70%; *P* = 3.789 × 10^−9^; Table S3 right) but not with shuffled data (45.8% - 54.7%; Table S4 right. We used an LDA to decode several levels together and found good visual assignment of the test bundles to the three revealed preference levels in our combined population of 24 participants (first discriminant; numeric accuracy of 88-100%; *P* = 9.46 × 10^−12^; Figure S4; three colored symbol groups) but not to different positions on same preference levels (second discriminant; numeric accuracy of 43-51%; *P* = 0.1433; mix of shapes) (note the reservations above when combining data from multiple participants). Thus, the two decoders showed together that the participants’ BDM bids distinguished bundles well between preference levels but not on the same preference level, thus confirming BDM validation of the two-dimensional revealed preference scheme of the ICs.

**Table S1A.** Test amounts (ml) of milkshake component A for stepwise psychophysical assessment of choice indifference points (IP): lowest indifference curve (IC1).

| Participant | Step 1 | Step 2 | Step 3 | Step 4 | Step 5 |
| --- | --- | --- | --- | --- | --- |
| 1 | 0.000 | 0.053 | 0.155 | 0.395 | 0.870 |
| 2 | 0.000 | 0.147 | 0.320 | 0.526 | 0.769 |
| 3 | 0.000 | 0.149 | 0.312 | 0.492 | 0.690 |
| 4 | 0.000 | 0.184 | 0.442 | 0.796 | 1.250 |
| 5 | 0.000 | 0.194 | 0.392 | 0.594 | 0.800 |
| 6 | 0.000 | 0.070 | 0.236 | 0.685 | 1.333 |
| 7 | 0.000 | 0.070 | 0.228 | 0.634 | 1.250 |
| 8 | 0.000 | 0.161 | 0.371 | 0.647 | 1.000 |
| 9 | 0.000 | 0.113 | 0.283 | 0.548 | 0.939 |
| 10 | 0.000 | 0.067 | 0.186 | 0.428 | 0.870 |
| 11 | 0.000 | 0.033 | 0.104 | 0.334 | 0.870 |
| 12 | 0.000 | 0.009 | 0.027 | 0.436 | 1.176 |
| 13 | 0.000 | 0.058 | 0.171 | 0.444 | 0.952 |
| 14 | 0.000 | 0.157 | 0.358 | 0.619 | 0.952 |
| 15 | 0.000 | 0.116 | 0.265 | 0.460 | 0.714 |
| 16 | 0.000 | 0.046 | 0.130 | 0.315 | 0.714 |
| 17 | 0.000 | 0.194 | 0.392 | 0.594 | 0.800 |
| 18 | 0.000 | 0.069 | 0.175 | 0.355 | 0.667 |
| 19 | 0.000 | 0.053 | 0.155 | 0.395 | 0.870 |
| 20 | 0.000 | 0.216 | 0.490 | 0.833 | 1.250 |
| 21 | 0.000 | 0.089 | 0.216 | 0.405 | 0.690 |
| 22 | 0.000 | 0.143 | 0.304 | 0.485 | 0.690 |
| 23 | 0.000 | 0.089 | 0.216 | 0.405 | 0.690 |
| 24 | 0.000 | 0.044 | 0.122 | 0.285 | 0.645 |
| MEAN | 0.000 | 0.105 | 0.252 | 0.504 | 0.894 |
| SEM | 0.000 | 0.012 | 0.024 | 0.030 | 0.044 |

Step 1: Reference Bundle set to (x = 0.0 ml; y = 2.0 ml). Steps 1- 5: test amounts of component A (x-axis) of the Variable Bundle for psychophysical variation of component B (y-axis).

**Table S1B.**
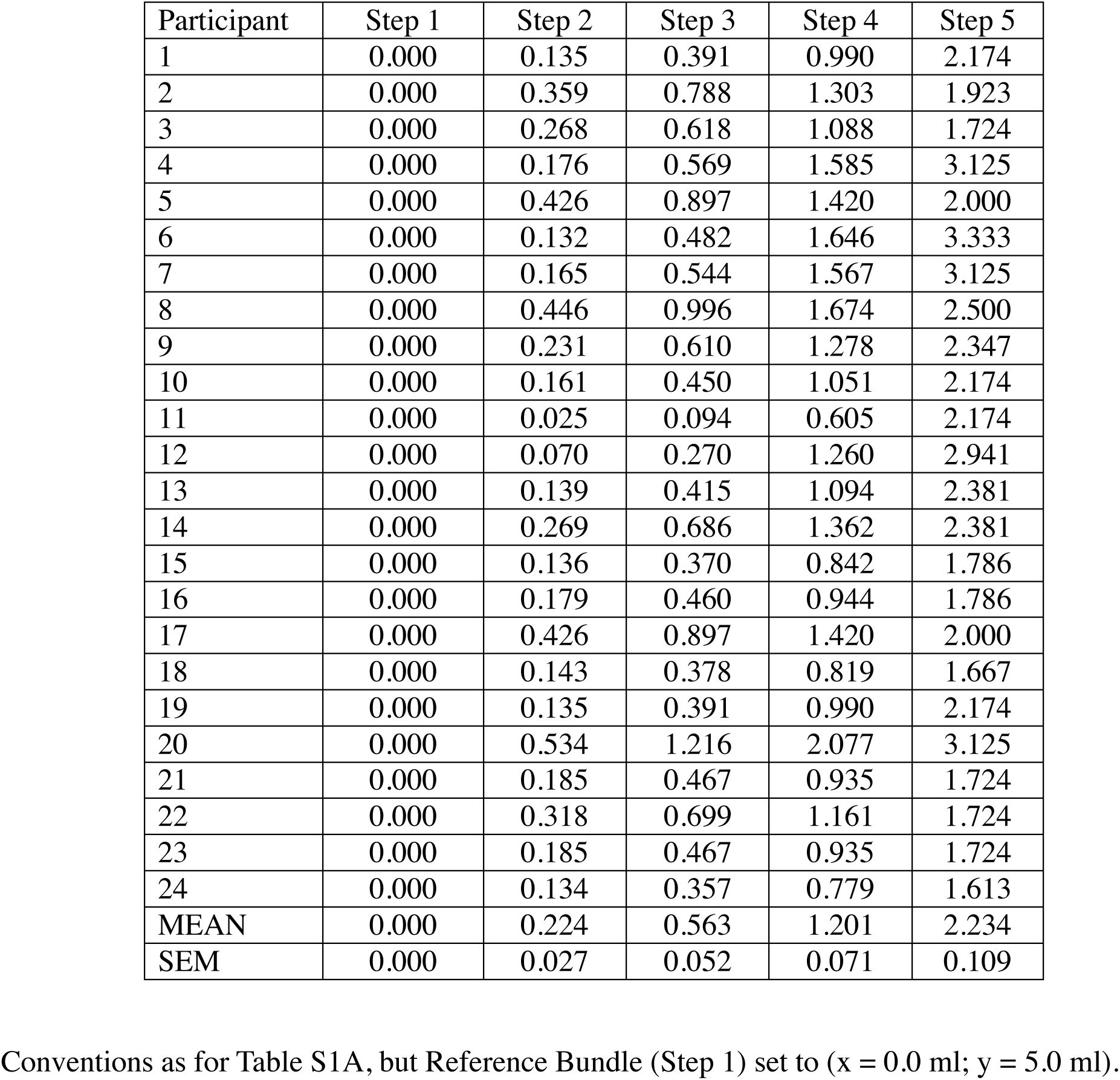
Test amounts (ml) of milkshake component A for stepwise psychophysical assessment of choice indifference points (IP): intermediate indifference curve (IC2).

**Table S1C.** Test amounts (ml) of milkshake component A for stepwise psychophysical assessment of choice indifference points (IP): highest indifference curve (IC3).

| Participant | Step 1 | Step 2 | Step 3 | Step 4 | Step 5 |
| --- | --- | --- | --- | --- | --- |
| 1 | 0.000 | 0.299 | 0.809 | 1.781 | 3.478 |
| 2 | 0.000 | 0.535 | 1.199 | 2.031 | 3.077 |
| 3 | 0.000 | 0.552 | 1.185 | 1.915 | 2.759 |
| 4 | 0.000 | 0.402 | 1.174 | 2.732 | 5.000 |
| 5 | 0.000 | 0.630 | 1.361 | 2.210 | 3.200 |
| 6 | 0.000 | 0.281 | 0.948 | 2.745 | 5.333 |
| 7 | 0.000 | 0.394 | 1.155 | 2.714 | 5.000 |
| 8 | 0.000 | 0.637 | 1.473 | 2.577 | 4.000 |
| 9 | 0.000 | 0.492 | 1.203 | 2.256 | 3.756 |
| 10 | 0.000 | 0.244 | 0.689 | 1.649 | 3.478 |
| 11 | 0.000 | 0.035 | 0.129 | 0.933 | 3.478 |
| 12 | 0.000 | 0.096 | 0.380 | 1.976 | 4.706 |
| 13 | 0.000 | 0.323 | 0.886 | 1.974 | 3.810 |
| 14 | 0.000 | 0.397 | 1.035 | 2.121 | 3.810 |
| 15 | 0.000 | 0.212 | 0.581 | 1.334 | 2.857 |
| 16 | 0.000 | 0.292 | 0.747 | 1.522 | 2.857 |
| 17 | 0.000 | 0.630 | 1.361 | 2.210 | 3.200 |
| 18 | 0.000 | 0.134 | 0.391 | 1.029 | 2.667 |
| 19 | 0.000 | 0.299 | 0.809 | 1.781 | 3.478 |
| 20 | 0.000 | 0.844 | 1.929 | 3.311 | 5.000 |
| 21 | 0.000 | 0.353 | 0.856 | 1.610 | 2.759 |
| 22 | 0.000 | 0.445 | 1.016 | 1.766 | 2.759 |
| 23 | 0.000 | 0.353 | 0.856 | 1.610 | 2.759 |
| 24 | 0.000 | 0.189 | 0.514 | 1.175 | 2.581 |
| MEAN | 0.000 | 0.378 | 0.945 | 1.957 | 3.575 |
| SEM | 0.000 | 0.039 | 0.081 | 0.119 | 0.175 |

Conventions as for Table S1A, but Reference Bundle (Step 1) set to (x = 0.0 ml; y = 8.0 ml).

**Table S2.** Amounts of milkshake component B (ml; y-axis) at the 7 test points for psychophysical estimation of indifference points for the lowest, intermediate and highest indifference curves (IC1 - IC3).

| Test point | 1 | 2 | 3 | 4 | 5 | 6 | 7 |
| --- | --- | --- | --- | --- | --- | --- | --- |
| IC1 | 0.000 | 0.333 | 0.667 | 1.000 | 1.333 | 1.667 | 2.000 |
| IC2 | 0.000 | 0.833 | 1.667 | 2.500 | 3.333 | 4.167 | 5.000 |
| IC3 | 0.000 | 1.333 | 2.667 | 4.000 | 5.333 | 6.667 | 8.000 |

**Table S3.** Accuracy (in %) of assigning bundles and BDM bids to original preference levels, using support vector machine decoder.

|  | Bundles |  |  | BDM bids |  |  |
| --- | --- | --- | --- | --- | --- | --- |
| Participant<br>C – convex IC<br>L – linear IC | Lev1<br>vs.<br>Lev2 | Lev1<br>vs.<br>Lev3 | Lev2<br>vs.<br>Lev3 | Lev1<br>vs.<br>Lev2 | Lev1<br>vs.<br>Lev3 | Lev2<br>vs.<br>Lev3 |
| C1 | 83.9 | 100.0 | 68.8 | 70.6 | 81.0 | 54.4 |
| C2 | 86.8 | 99.8 | 68.7 | 44.0 | 51.0 | 47.4 |
| C3 | 72.7 | 96.0 | 58.7 | 56.6 | 71.3 | 59.3 |
| C4 | 88.0 | 99.5 | 67.6 | 67.7 | 76.3 | 62.5 |
| C5 | 99.1 | 100.0 | 81.4 | 64.8 | 69.9 | 49.8 |
| C6 | 93.7 | 99.5 | 61.9 | 53.5 | 65.9 | 54.0 |
| C7 | 66.2 | 75.5 | 49.5 | 70.0 | 72.2 | 43.6 |
| C8 | 72.9 | 81.8 | 52.6 | 49.8 | 67.2 | 62.6 |
| C9 | 80.0 | 100.0 | 80.0 | 68.4 | 83.8 | 62.1 |
| C10 | 99.6 | 100.0 | 69.8 | 47.0 | 46.6 | 43.7 |
| C11 | 90.2 | 99.5 | 55.4 | 68.9 | 75.7 | 53.1 |
| C12 | 96.8 | 100.0 | 71.2 | 47.3 | 57.9 | 47.1 |
| C12 | 86.1 | 87.7 | 56.7 | 58.7 | 60.9 | 48.2 |
| C14 | 80.0 | 100.0 | 80.0 | 63.2 | 74.3 | 56.8 |
| C15 | 99.8 | 100.0 | 85.5 | 67.8 | 78.6 | 54.8 |
| C16 | 100.0 | 100.0 | 90.0 | 57.8 | 62.5 | 50.4 |
| C17 | 100.0 | 100.0 | 100.0 | 44.8 | 70.9 | 68.4 |
| C18 | 99.0 | 99.9 | 55.3 | 68.1 | 75.3 | 51.7 |
| L19 | 100.0 | 100.0 | 88.6 | 43.5 | 65.8 | 59.6 |
| L20 | 100.0 | 100.0 | 95.9 | 69.2 | 84.9 | 70.1 |
| L21 | 100.0 | 100.0 | 100.0 | 62.2 | 66.0 | 44.4 |
| L22 | 100.0 | 100.0 | 83.6 | 54.8 | 52.5 | 45.9 |
| L23 | 100.0 | 100.0 | 100.0 | 63.2 | 58.4 | 47.8 |
| L24 | 100.0 | 100.0 | 80.0 | 61.4 | 68.5 | 57.7 |

Accuracy is shown in % of numbers of bundles and BDM bids correctly assigned to one of the two tested revealed preference levels (averages from 150 iterations of classification of 10 pseudorandomly selected bundles / BDM bids from two tested revealed preference levels). Bundles were tested at psychophysically assessed choice indifference points (IP) on one of three revealed preference levels (corresponding to the three indifference curves, IC, fitted to the IPs). The tested bundles had been pseudorandomly selected from 5 unique estimated IPs on each of two revealed preference levels under study. BDM: Becker-DeGroot-Marschak auction-like bidding mechanism. Lev1, Lev2, Lev3: revealed preference levels, numbered according to distance from origin. The 24 participants were labelled as C or L according to their IPs being fitted best to convex or linear ICs.

**Table S4.** Chance decoding accuracy of shuffled data by support vector machine.

| Participant<br>C – convex IC<br>L – linear IC | Bundles |  |  | BDM bids |  |  |
| --- | --- | --- | --- | --- | --- | --- |
|  | Lev1<br>vs.<br>Lev2 | Lev1<br>vs.<br>Lev3 | Lev2<br>vs.<br>Lev3 | Lev1<br>vs.<br>Lev2 | Lev1<br>vs.<br>Lev3 | Lev2<br>vs.<br>Lev3 |
| C1 | 53.5 | 50.9 | 51.3 | 45.8 | 49.1 | 52.0 |
| C2 | 49.6 | 48.9 | 50.0 | 50.1 | 51.7 | 53.2 |
| C3 | 50.2 | 52.5 | 51.0 | 50.3 | 50.2 | 50.8 |
| C4 | 48.6 | 48.2 | 52.0 | 53.1 | 50.2 | 50.1 |
| C5 | 48.8 | 51.8 | 51.2 | 51.6 | 50.7 | 50.0 |
| C6 | 46.8 | 50.3 | 53.2 | 48.4 | 49.2 | 50.2 |
| C7 | 48.3 | 51.3 | 49.2 | 49.7 | 50.2 | 49.7 |
| C8 | 48.9 | 48.9 | 49.2 | 50.2 | 51.6 | 50.1 |
| C9 | 50.3 | 52.0 | 46.9 | 52.0 | 50.0 | 49.9 |
| C10 | 49.7 | 52.4 | 49.4 | 47.2 | 53.0 | 51.4 |
| C11 | 48.6 | 52.2 | 44.7 | 49.2 | 50.4 | 49.6 |
| C12 | 49.5 | 53.1 | 46.9 | 52.3 | 48.7 | 50.5 |
| C12 | 49.6 | 48.9 | 49.5 | 50.3 | 53.1 | 50.3 |
| C14 | 52.1 | 51.7 | 51.3 | 52.3 | 52.0 | 50.4 |
| C15 | 49.2 | 53.0 | 47.2 | 52.4 | 49.5 | 47.7 |
| C16 | 53.4 | 52.5 | 47.2 | 46.5 | 51.1 | 47.1 |
| C17 | 49.8 | 53.0 | 50.7 | 52.9 | 52.0 | 51.0 |
| C18 | 49.8 | 48.2 | 52.0 | 51.4 | 48.4 | 49.9 |
| L19 | 48.4 | 53.7 | 52.6 | 50.0 | 50.7 | 47.0 |
| L20 | 50.0 | 53.4 | 49.8 | 49.7 | 54.1 | 52.2 |
| L21 | 54.6 | 53.4 | 48.5 | 54.2 | 52.4 | 54.7 |
| L22 | 47.0 | 52.7 | 51.1 | 51.9 | 48.9 | 47.7 |
| L23 | 51.4 | 53.1 | 47.2 | 50.4 | 51.2 | 50.1 |
| L24 | 49.3 | 50.3 | 48.6 | 49.7 | 49.9 | 49.5 |

The support vector machine analysis used the shuffled matrix of 2 columns (containing the x- and y-coordinates of bundle components A and B, respectively) and 5 rows (containing the 5 bundles). Conventions as for Table S4.

**Table S5.** Two-way Analysis of Variance (ANOVA) of BDM bids.

| Participant | 1st factor F(2,179) |  |  | 2nd factor F(4,179) |  |  | Interaction F(8,179) |  |  |
| --- | --- | --- | --- | --- | --- | --- | --- | --- | --- |
|  | P | F | MSE | P | F | MSE | P | F | MSE |
| 1* | $9.98 \times 10^{-46}$ | 128.68 | 1656.30 | 0.01* | 2.07 | 26.70 | 0.99 | 0.45 | 5.78 |
| 2 | $5.37 \times 10^{-40}$ | 109.15 | 1829.07 | 0.53 | 0.93 | 15.51 | 1.00 | 0.34 | 5.74 |
| 3 | $5.63 \times 10^{-63}$ | 194.16 | 2148.07 | 0.61 | 0.86 | 9.47 | 0.70 | 0.84 | 9.32 |
| 4* | $5.68 \times 10^{-79}$ | 265.03 | 2585.34 | 0.02* | 1.99 | 19.37 | 0.90 | 0.67 | 6.50 |
| 5* | $3.53 \times 10^{-56}$ | 167.10 | 2300.07 | 0.03* | 1.84 | 25.28 | 0.71 | 0.84 | 11.51 |
| 6 | $2.13 \times 10^{-49}$ | 141.75 | 1773.60 | 0.64 | 0.83 | 10.39 | 0.96 | 0.58 | 7.30 |
| 7 | $8.67 \times 10^{-41}$ | 111.79 | 1620.12 | 0.83 | 0.64 | 9.28 | 1.00 | 0.20 | 2.85 |
| 8 | $4.25 \times 10^{-58}$ | 174.57 | 1793.40 | 0.97 | 0.41 | 4.24 | 1.00 | 0.21 | 2.18 |
| 9 | $8.04 \times 10^{-60}$ | 181.39 | 1716.27 | 0.98 | 0.39 | 3.65 | 1.00 | 0.14 | 1.30 |
| 10 | $9.28 \times 10^{-34}$ | 89.04 | 1656.30 | 0.80 | 0.68 | 12.63 | 1.00 | 0.27 | 5.10 |
| 11 | $9.81 \times 10^{-53}$ | 154.02 | 1681.34 | 0.88 | 0.58 | 6.31 | 1.00 | 0.26 | 2.79 |
| 12 | $5.38 \times 10^{-116}$ | 475.79 | 4001.40 | 0.98 | 0.38 | 3.22 | 1.00 | 0.21 | 1.78 |
| 13 | $6.88 \times 10^{-49}$ | 139.91 | 1956.07 | 0.38 | 1.08 | 15.07 | 0.99 | 0.46 | 6.37 |
| 14 | $1.27 \times 10^{-65}$ | 205.17 | 2160.00 | 0.93 | 0.51 | 5.39 | 1.00 | 0.19 | 2.04 |
| 15 | $5.86 \times 10^{-63}$ | 194.09 | 2527.41 | 0.79 | 0.69 | 8.97 | 1.00 | 0.34 | 4.37 |
| 16 | $7.10 \times 10^{-52}$ | 150.82 | 2383.27 | 0.77 | 0.70 | 11.09 | 0.99 | 0.49 | 7.68 |
| 17 | $1.82 \times 10^{-74}$ | 243.99 | 2574.67 | 0.69 | 0.78 | 8.20 | 1.00 | 0.19 | 1.99 |
| 18 | $2.81 \times 10^{-70}$ | 225.21 | 2667.23 | 0.68 | 0.79 | 9.40 | 0.99 | 0.48 | 5.72 |
| 19 | $2.13 \times 10^{-48}$ | 138.15 | 2005.07 | 0.83 | 0.64 | 9.27 | 0.99 | 0.45 | 6.52 |
| 20 | $4.82 \times 10^{-58}$ | 174.35 | 1865.36 | 0.87 | 0.59 | 6.32 | 1.00 | 0.25 | 2.69 |
| 21 | $1.61 \times 10^{-72}$ | 235.16 | 2486.40 | 0.43 | 1.02 | 10.82 | 1.00 | 0.28 | 2.99 |
| 22 | $2.67 \times 10^{-56}$ | 167.56 | 1658.40 | 0.07 | 1.63 | 16.15 | 0.60 | 0.91 | 8.98 |
| 23 | $4.61 \times 10^{-108}$ | 424.33 | 4768.92 | 0.05 | 1.70 | 19.15 | 0.37 | 1.07 | 12.07 |
| 24 | $1.49 \times 10^{-73}$ | 239.83 | 2029.07 | 0.39 | 1.06 | 9.00 | 0.40 | 1.05 | 8.89 |

F(df1, df2): 1st factor: df1 = 2 (n - 1, n = 3 preference levels), df2 = 179 (n - 1, n = 5 IPs x 3 levels x 12 trial repetitions = 180); 2nd factor: df1 = 4 (n - 1, n = 5 IPs per preference level), df2 = 179; Interaction: df1 = 8 ((n - 1) x (m - 1), n = 3 preference levels, m = 5 IPs per preference level), df2 = 179. df: degree of freedom; IP: bundle at indifference point; MSE: mean square error; \* Exceptional 3 participants with significantly different BDM bids in 2nd factor ( $P < 0.05$ ).

**Figure S1.**
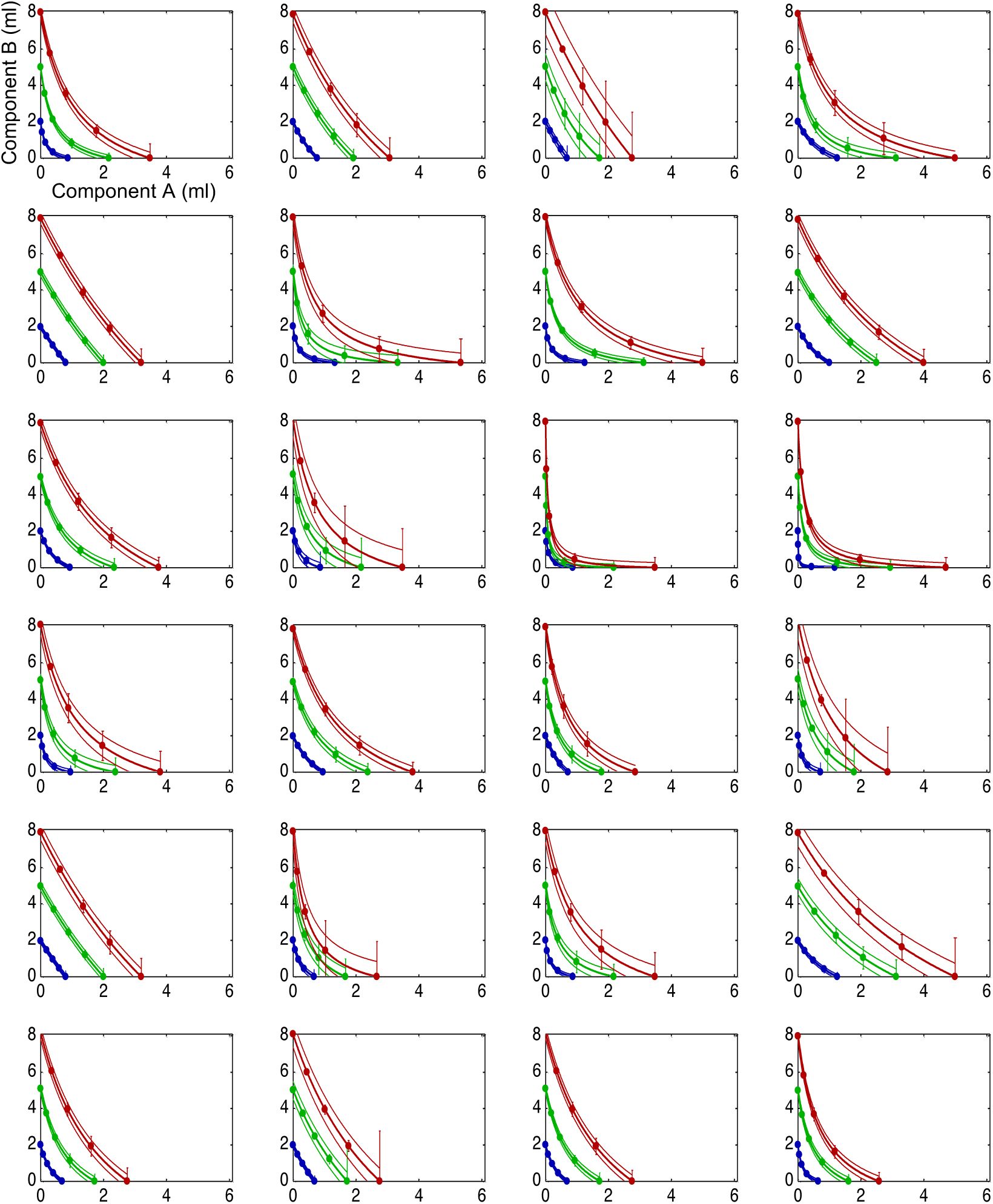
Empirical indifference curves (IC) from all 24 participants. Note the fanning out of the confidence intervals towards the bottom right in each graph (thin lines), which likely reflects the progression of choice testing: the Reference Bundle was kept constant at the y-axis intercept (x = 0), whereas testing with the Variable Bundle progressed from top left to bottom right. Same conventions as for Figure 2.

**Figure S2.**
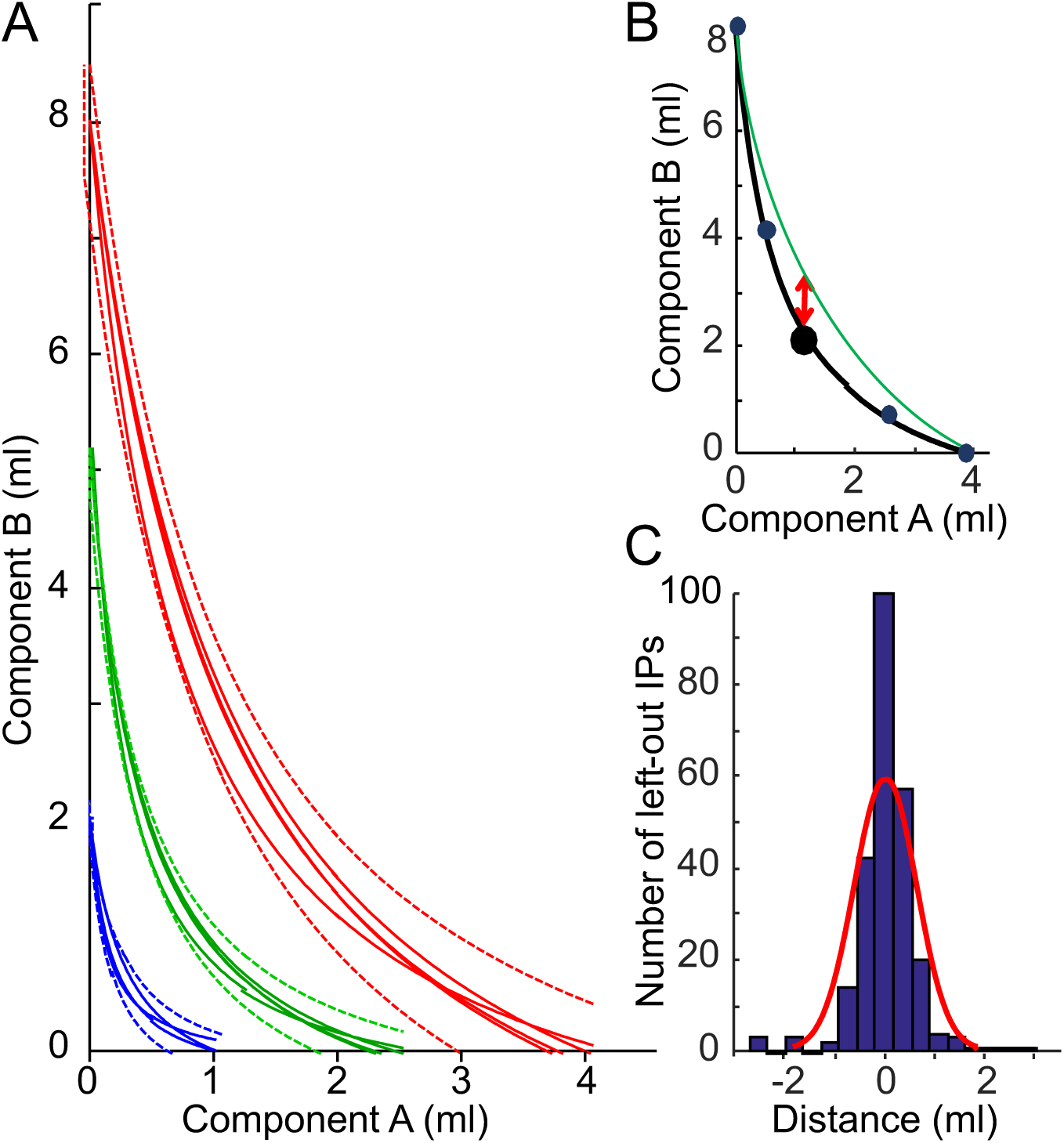
Leave-one-out validation of estimated indifference curves (ICs). (A) Graphic assessment with an example participant: hyperbolically refitted ICs with one left-out indifference point (IP) (solid lines), plotted together with 95% confidence intervals of the original hyperbolically fitted ICs (dotted lines). The refitting resulted in 4 new ICs (partly overlapping) at each of three levels. None of the refitted IPs fell outside the original confidence intervals. (B) Scheme of numeric assessment: distance in ml (in ml on y-axis, red) between the left-out IP on the original IC (heavy black dot on black curve) and the refitted IC (green). (C) Histogram of distance between refitted ICs and left-out IPs across all subjects. Skewness was - 0.12, suggesting rather symmetric distribution around the mean (modeled in red).

**Figure S3.**
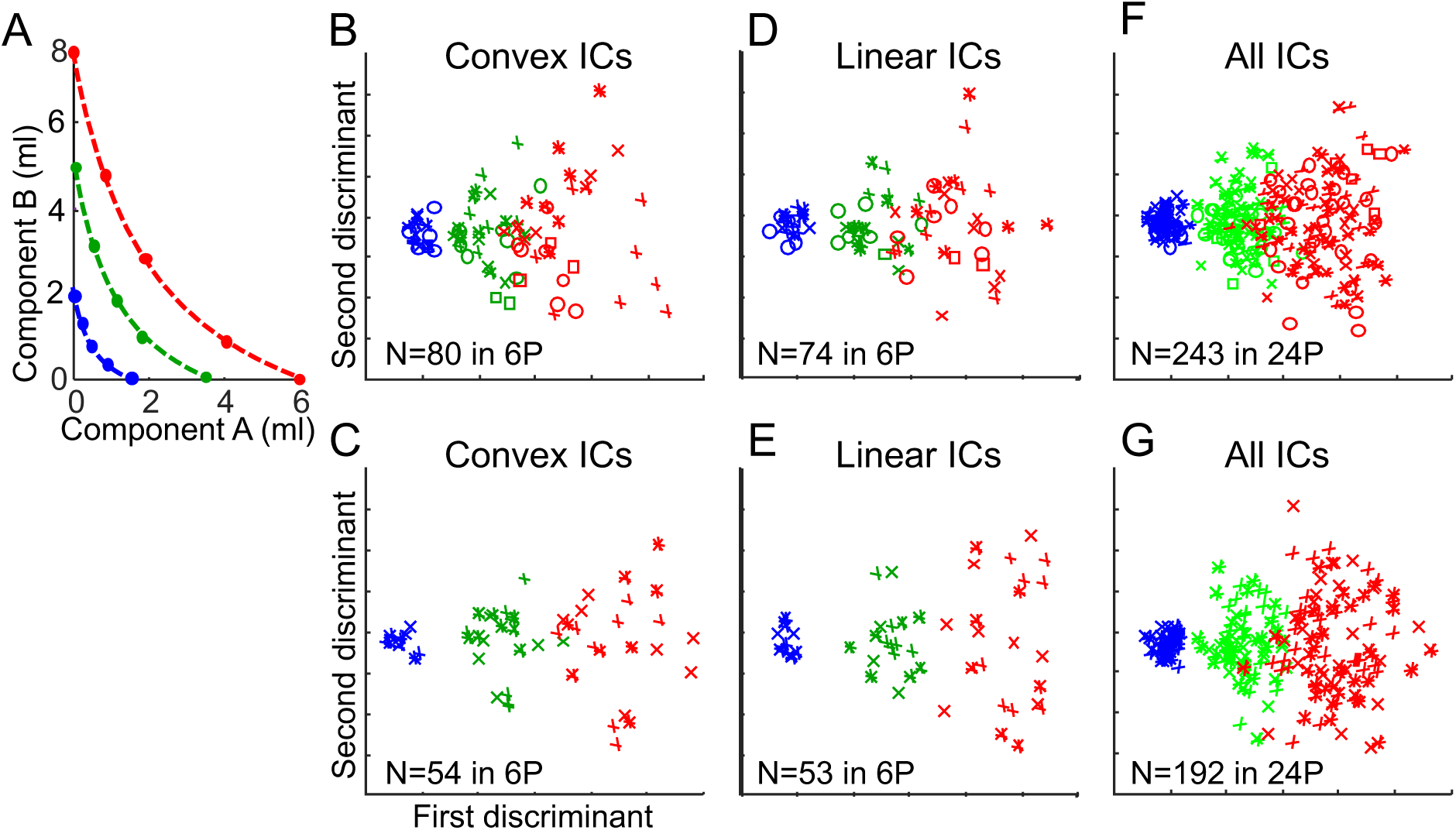
Visualization of bundle decoding using Linear Discriminant Analysis (LDA). (A) Schematics of bundle decoding at psychophysically estimated points of equal revealed preference (indifference points, IPs, plotted along the dotted lines). Following the notions of revealed preference, LDA should show accurate decoding between the three preference levels (green, blue, red) but not along each level. (B) LDA bundle distinction between three levels of revealed preference (81-100% binary numeric decoding accuracy between two levels; first discriminant) but not along same preference levels (second discriminant) in N = 80 bundles from six participants (6P) with similar convex ICs (same convention applies to all panels). Bundles on the three preference levels are colored blue, green and red according to distance from origin, red being highest. The five bundles on each preference level are marked from top left to bottom right with ‘o’, ‘*’, ‘+’, ‘x’ and ‘□’ symbols. Due to the arbitrariness of the scale, numbers are not indicated. (C) As B but for partly physically non-dominating bundles (one lower component in preferred bundle than in alternative bundle (97-100% accuracy; first discriminant). (D) As B but for six participants with linear ICs (90-100% accuracy). (E) As D but for partly physically non-dominating bundles (96-100% accuracy). (F) As B but for all 24 participants (18 with convex IC, six with linear ICs) (83-100% accuracy). (G) As F but for partly physically non-dominating bundles (97-100% accuracy).

**Figure 4.**
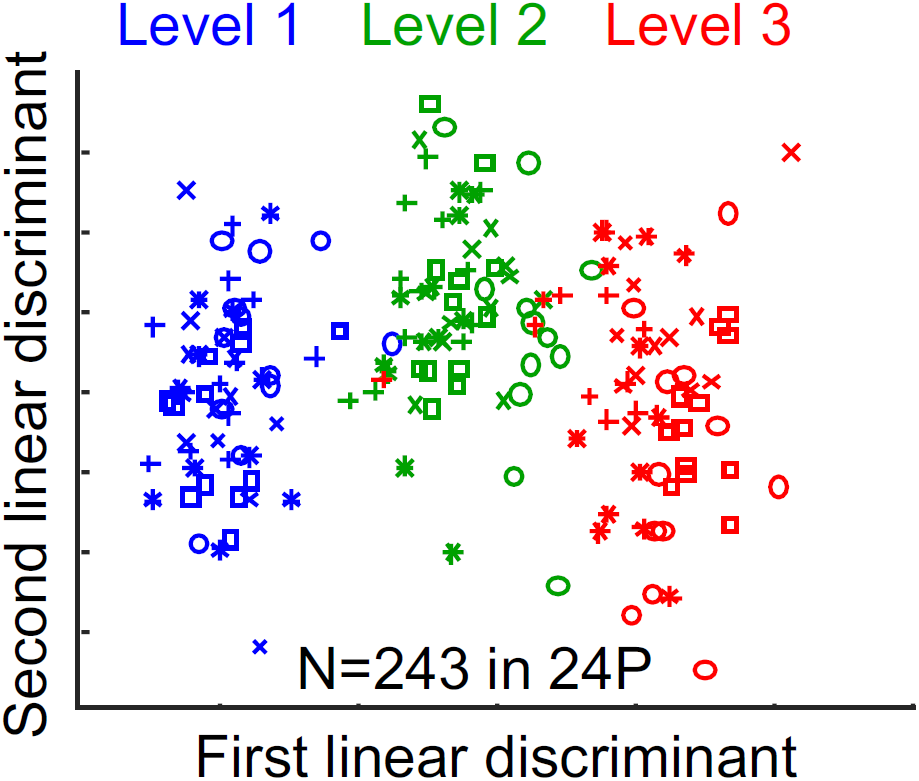
Visual decoding by Linear Discriminant Analysis (LDA) of BDM bids for bundles on different revealed preference levels (first discriminant; 88-100% numeric decoding accuracy; *P* = 9.46 × 10^−12^) and along same preference levels (second discriminant; 43-51% accuracy) (N = 243 bundles; all 24 participants). The LDA used scalar BDM bids for bundles positioned at IPs shown in Figure 4A.

